# Neutrophil NADPH oxidase breaks the inflammatory IL-1β/IL-17A circuit to enhance pathogen clearance during respiratory virus infections

**DOI:** 10.1101/2024.09.12.612505

**Authors:** Aderonke Sofoluwe, Angelos Petropoulos, Abhilesh Salil Goomanee, Zehra Fatima Ali-Khan, Annika Warnatsch

## Abstract

Respiratory virus infections are invariably accompanied by an increase in oxidative stress, characterised by elevated production of Reactive Oxygen Species (ROS) in the lung, which plays a pivotal role in both pathogenesis and host defence. Using a mouse model, neutrophil NADPH Oxidase 2 (Nox2) emerges as a key player, primarily responsible for generation of ROS during the early phases of Influenza A Virus (IAV) infection. Neutrophil Nox2-derived ROS display a multifaceted role, not only unleashing oxidative stress but in turn curbing neutrophil-derived IL-1β signalling. Absence of neutrophil Nox2 triggered heightened production of IL-1β, promoting the proliferation of IL-17-producing gamma delta (γδ) T cells. This early self-amplified augmentation of the IL-β/IL-17 axis counteracted the antiviral interferon response against IAV infection in mice. We extended our findings to humans. Similar patterns of ROS production and cytokine regulation were observed in human neutrophils when exposed to virus analogue poly(I:C) and SARS-CoV-2. Our discovery highlights that ROS, often associated with harm, play a dual role by regulating cytokine signalling and thus influencing the immune response against respiratory viruses.

**Highlights:**

- Neutrophil NADPH oxidase 2 is the predominant source of ROS production early after influenza A virus infection.
- Neutrophil Nox2-derived ROS uniquely suppress IL-1β, influencing γδ^17^ T cell proliferation.
- In absence of neutrophil Nox2, increased neutrophil-derived IL-1β drives proliferation of γδ^17^ T cells.
- Early amplification of the IL-β/IL-17 axis counteracts the antiviral interferon response in IAV-infected mice.
- Virus infection induces a self-perpetuating loop between human blood-derived IL-1β-producing neutrophils and IL-17-secreting γδ T cells.

## Introduction

Respiratory viral infections such as Influenza A Virus (IAV) and SARS-CoV-2 represent a major global health challenge due to their high prevalence and substantial impact on public health. The innate immune system plays a crucial frontline role in the body’s defence against respiratory viruses.^(1)^ In the respiratory tract, the primary entry point for viruses, epithelial cells and resident immune cells like alveolar macrophages and dendritic cells detect virus presence through pattern recognition receptors (PRRs) such as toll-like receptors (TLRs) and RIG-I-like receptors. This triggers a signalling cascade that produces type I interferons (IFNs), cytokines and chemokines. Type I IFNs are critical for inducing an antiviral state by activating IFN-stimulated genes, thereby inhibiting viral replication and spread. Concurrently, cytokines and chemokines facilitate inflammation and activate the adaptive immune response while recruiting additional immune cells, such as neutrophils and NK cells, to the infection site.

Neutrophils, one of the first responders, engage in phagocytosis, release neutrophil extracellular traps (NETs), and produce reactive oxygen species (ROS) to contain viral spread.^(2)^ Depletion studies in mice have shown important roles for neutrophils in controlling viral replication and directing subsequent immune responses.^(3–5)^ However, their overactivation can lead to pulmonary inflammation and damage, reflecting the immune system’s need to balance effective viral defence with potential tissue injury.^(6,7)^

The induction of oxidative stress via increased ROS production is a critical component of the pathophysiology of respiratory virus infections. Understanding the dual nature of ROS in these infections is crucial, as they are not only damaging agents but also play pivotal roles in immune signalling, cytokine production, inflammation, and cell death.^(8)^ Production of radicals and redox imbalance increase with age^(9)^ and so does susceptibility to fatal outcomes to virus infection such as SARS-CoV-2. This complex interplay highlights the importance of understanding both the detrimental and beneficial effects of ROS, particularly in the context of viral replication and associated inflammation.

The neutrophil NADPH oxidase 2 (Nox2) is a multi-subunit enzyme complex primarily responsible for rapid ROS production during the oxidative burst.^(10)^ This mechanism, while essential for pathogen killing, must be carefully regulated. Besides their microbicidal functions, Nox2-derived ROS act as signalling molecules with the capacity to regulate cellular pathways through reduction-oxidation (redox) regulation of target proteins.^(11)^ Thus, the balance of Nox2 activity is crucial, with its dysregulation being implicated in severe disease outcomes in respiratory viral infections such as Respiratory Syncytial Virus (RSV)^(12)^, IAV^(13)^ and SARS-CoV-2.^(14)^

Studies in mice with complete knockout of Nox2 have demonstrated improved clearance of lung influenza infection with increased cytokine levels, concluding that absence of Nox2 improved the resolution of IAV infection but not inflammation.^(15, 16)^ To et al. showed that endosomal ROS production in macrophages in response to viral infection causes oxidation of a highly conserved cysteine residue in Toll-like receptor 7 (TLR7), abrogating TLR7 activation and ultimately suppressing antiviral interferon IFNβ but also production of IL-1β and TNFα.^(17)^ However, the amount of ROS produced by macrophages is negligible compared to that of neutrophil-derived ROS, making neutrophil ROS production during respiratory virus infection a crucial topic.

Previous studies have implicated ROS in the control of inflammatory responses, underlying the dual role of neutrophil-derived ROS in infection and inflammation, such as through the inhibition of cytokine production.^(18–22)^ Evidence for the anti-inflammatory role of Nox2-derived ROS has been demonstrated in various disease models via different mechanisms. For example, Nox2-deficient mice in a model of lipopolysaccharide-induced lung inflammation display higher proinflammatory cytokine levels and increased infiltration of neutrophils in the lung, controlled via redox regulation of transcription factor NFκB ^(18)^ and caspase-1.^(19)^ Further studies have demonstrated redox sensitivity of NFκB in neutrophils regulating immune responses to bacterial and fungal lung infection.^(20)^ Moreover, ROS-deficient neutrophils have been shown to overproduce IL-8^(21)^, indicating their capacity to act as regulatory cells inhibiting inflammatory circuits.

Neutrophil Nox2-mediated ROS production has been shown to directly impact cytokine production in adjacent immune cells via the transfer of extracellular ROS. In a human sepsis model, neutrophils have been shown to inhibit systemic inflammation by ROS-dependent suppression of CD8 T cells in a contact-dependent manner.^(23)^ Additionally, human and mouse neutrophils have been suggested to suppress proinflammatory gamma delta (γδ) T cell responses in different models.^(24–26)^

Despite these insights, there is limited understanding of the specific roles of neutrophil-derived ROS in modulating the immune response during respiratory viral infections. Our study addresses this gap by exploring the dual role of ROS, produced by Nox2, in suppressing IL-1β and IL-17A signalling. Understanding this regulatory mechanism is essential because IL-1β and IL-17A are pivotal in driving inflammatory responses, and their dysregulation can lead to severe tissue damage and impaired viral clearance. The balance between these pro-inflammatory cytokines and antiviral interferon responses is crucial for effective immunity.

Moreover, previous studies have shown the overall impact of ROS on immune responses, but there has been limited understanding of cell-specific ROS functions. Our study specifically focuses on neutrophil-derived ROS and their impact on other immune cells, particularly γδ T cells.

Using advanced mouse models and extending to human cells, we found that neutrophil NADPH Oxidase regulates antiviral immune responses by modulating IL-1β and IL-17A signalling. Our results highlight neutrophil Nox2 as the predominant source of ROS during early phases of respiratory virus infections and its unique role in modulating inflammatory signalling pathways. We demonstrate how neutrophil-derived ROS affect γδ T cell proliferation and function, which has implications for understanding the broader immune landscape during viral infections. In absence of neutrophil Nox2, increased neutrophil-derived IL-1β drives proliferation of γδ^17^ T cells. This early amplification of the IL-β/IL-17 axis counteracts the antiviral interferon response in IAV-infected mice. Additionally, viral infection induced a self-perpetuating loop between human blood-derived IL-1β-producing neutrophils and IL-17-secreting γδ T cells *ex vivo*. By elucidating the dual role of ROS, our results suggest potential therapeutic strategies that involve modulating ROS levels to balance antiviral immunity and inflammation.

## Results

### Neutrophil Nox2 is a predominant source of ROS early after IAV lung infection

To study the effects of the neutrophil oxidative burst without affecting ROS production in other cells, we generated *Ly6G-Cre(+/−)*/tdTomato *Nox2 flox* mice using the neutrophil-specific *Ly6G* locus. *Ly6G-Cre* mice provide to-date the most specific approach to study and track neutrophils.^(27)^ We firstly confirmed Nox2 deletion in neutrophils of *Ly6G-Cre(+/−)*/tdTomato *Nox2 flox* mice by immunofluorescence staining for Nox2 subunit gp91phox in lungs (Figure S1A) and flow cytometry on blood (Figure S1B and C). Neutrophils (gated as Live CD45^+^, CD11b^+^ and Ly6G^+^ cells) of *Ly6G-Cre(+/−)*/tdTomato *Nox2 flox* mice expressed tdTomato and lacked Nox2 expression.

To analyse the contribution of neutrophil Nox2-mediated superoxide production to overall ROS levels in IAV-infected lungs, we infected *Ly6G-Cre*(+/−) *Nox2 flox* mice with IAV strain X-31 at 3×10^4^ TCID_50_ – a mild and self-resolving infectious dose - and measured superoxide production in the lung by Dihydroethidium (DHE) staining 18 hours post-infection (Figure 1A and B). Compared to IAV-infected *Ly6G-Cre(-/-) Nox2 flox* control mice DHE Staining was 1.8-fold reduced in *Ly6G-Cre(+/−) Nox2 flox* mice comparable to *Nox2-/-* mice. Moreover, we assessed hydrogen peroxide levels by CellROX Green staining. Percentage of Live CD45^+^ CellROX^+^ immune cells in the lungs of IAV-infected *Ly6G-Cre(-/-) Nox2 flox* mice increased to 38.2 ± 18.7 % at day 3 post-infection while in *Ly6G-Cre(+/−) Nox2 flox* mice only 12.4 ± 4.3 % of lung immune cells were CellROX^+^ comparable to mock-infected control and IAV-infected *Nox2-/-* mice (13.4 ± 2.9 % and 10.4 ± 4.2 %, respectively; Figure 1C). The CD45^-^ non-immune cell compartment showed unimpaired induction of oxidative stress in *Ly6G-Cre(+/−) Nox2 flox* mice (Figure 1D). Baseline CellROX^+^ intensity in the CD45^-^ compartment of Nox2-/- mice was elevated which might be attributed to compensatory ROS production via Nox2-independent pathways that lead to direct production of hydrogen peroxide, however no upregulation was found post-IAV infection. The similarity between neutrophil-specific Nox2-deficient and *Nox2-/-* animals revealed that neutrophil Nox2 is the predominant source of ROS in the immune cell compartment between 1 to 3 days post-IAV infection. While ROS production in the non-immune compartment remained intact.

**Figure 1.**
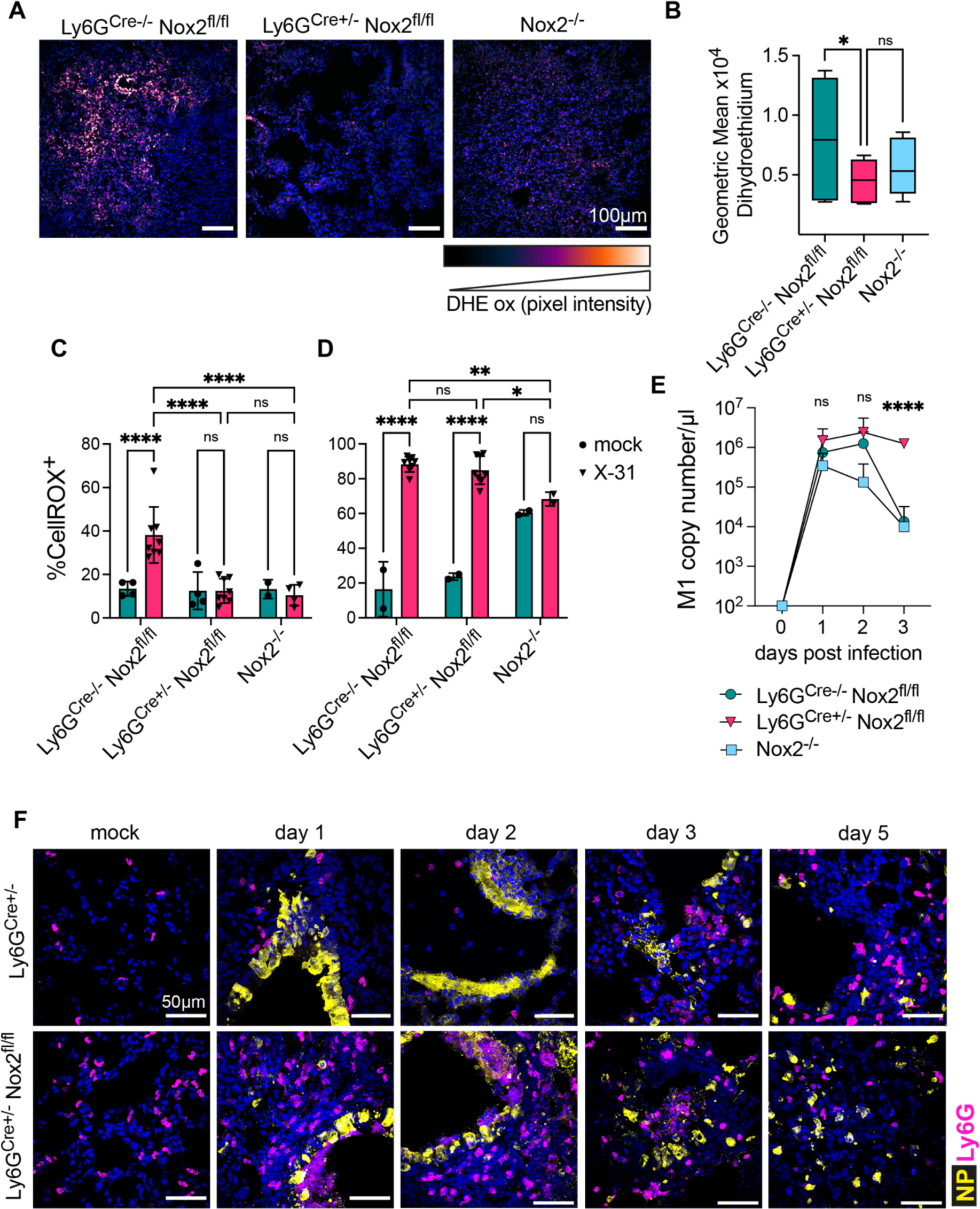
Neutrophil Nox2-derived ROS play crucial role in viral clearance during early IAV infection. *Ly6G-Cre*^-/-^ *Nox2^fl/fl^*, *Ly6G-Cre*^+/−^ *Nox2^fl/fl^* and *Nox2^-/-^* mice were infected with IAV strain X-31 at 3×10^4^ TCID_50_ or PBS (mock) in 30 µl intranasally. A - B) Representative micrographs of lung sections stained with Dihydroethidium (DHE) for superoxide 18 hrs after infection (A) and DHE geometric mean of lung lymphocytes (B) (n = 3). C - D) Percentage of CellROX^+^ cells gated as Live CD45^+^ (C) and Live CD45^-^ (D) in the lung 3 days post-infection (n = 2 - 4 mock, n = 8 X-31 for *Ly6G-Cre*^-/-^ *Nox2^fl/fl^* and *Ly6G-Cre*^+/−^ *Nox2^fl/fl^,* n = 2 – 4 for *Nox2^-/-^*). E) RT-qPCR of influenza matrix protein M1 in whole lungs at indicated time points post-infection (n = 6 - 8). F) Representative micrographs of lung sections stained for IAV nucleoprotein (NP, yellow), neutrophils (Ly6G, magenta) and nuclei (DAPI, blue) at indicated time points post-infection. Scale bar 50 µm. Bars represent mean ± SEM of 2 – 3 independent experiments. p values are indicated, *<0.05, **<0.01, ***<0.001, ns not significant. p values were determined using two-way ANOVA and Tukey’s post-test.

### Absence of neutrophil Nox2 impairs virus clearance and increases lung inflammation

Previous work has demonstrated improved resolution of IAV infection in *Nox2-/-* mice. ^(15,16)^ To examine how neutrophil Nox2-derived ROS production affects virus replication we measured viral matrix mRNA in the lung after infection with IAV strain X-31 at 3×10^4^ TCID_50_ (Figure 1E and S2A). After initial increase of viral burden 48 hours post-infection in all mouse strains, viral burden rapidly declined in *Nox2-/-* as well as *Ly6G-Cre(-/-) Nox2 flox* control mice at day 3 post-infection, while viral burden did not significantly reduce in *Ly6G-Cre(+/−) Nox2 flox* mice. We confirmed diminished virus clearance in *Ly6G-Cre(+/−) Nox2 flox* mice by immunofluorescence staining for viral nucleoprotein (NP, yellow) in lung sections of infected mice (Figure 1F). Interestingly, increased accumulation of neutrophils (Ly6G, magenta) was observed around infected bronchioli and larger amounts of viral nucleoprotein persisted in infected lungs of *Ly6G-Cre(+/−) Nox2 flox* mice until day 5 post-infection. Reduced virus clearance was further corroborated by significantly increased weight loss in IAV-infected *Ly6G-Cre(+/−) Nox2 flox* mice (Figure S2B). Histopathological analysis revealed a transient increase in inflammation in *Ly6G-Cre(+/−) Nox2 flox* mice from day 1 to day 3 compared to *Ly6G-Cre(-/-) Nox2 flox* control mice as measured by nuclei count and scoring (Figures S2C, D and E). Inflammation in *Ly6G-Cre(-/-) Nox2 flox* mice concentrated around bronchioli. In contrast, lung damage spread further to alveoli in *Ly6G-Cre(+/−) Nox2 flox* mice. Overall, neutrophil-specific Nox2-deficient mice exhibit heightened lung inflammation and damage, coupled with compromised virus clearance.

### The immune response to influenza A virus is shifted towards IL-1β/IL-17A axis in neutrophil-specific Nox2-deficient mice

The composition of cellular infiltrates in lung tissue was investigated by flow cytometry. Total lung cell counts were 1.5-fold increased in *Ly6G-Cre(+/−) Nox2 flox* compared to *Ly6G-Cre(+/−)* control mice 3 days post-infection (Figure 2A). Predominantly, numbers of neutrophils and T cells were increased in IAV-infected *Ly6G-Cre(+/−) Nox2 flox* mice (Figure 2B and D). We further observed elevated counts of inflammatory Ly6C^hi^ monocytes (Figure 2C). Interestingly, CD8^+^ and CD4^+^ αβ T cells cannot alone account for the increased number of CD3^+^ T cells (Figure 2E). No changes were observed in cell counts of macrophages, alveolar macrophages, conventional dendritic cells or NK cells (Figure S3A, B, C and D). No significant differences in lung immune cell counts between *Ly6G-Cre*(+/−) *Nox2 flox and Ly6G-Cre(+/−)* control mice were observed 7 days post-IAV infection and total lung cell counts declined in both strains (Figure 2A).

**Figure 2.**
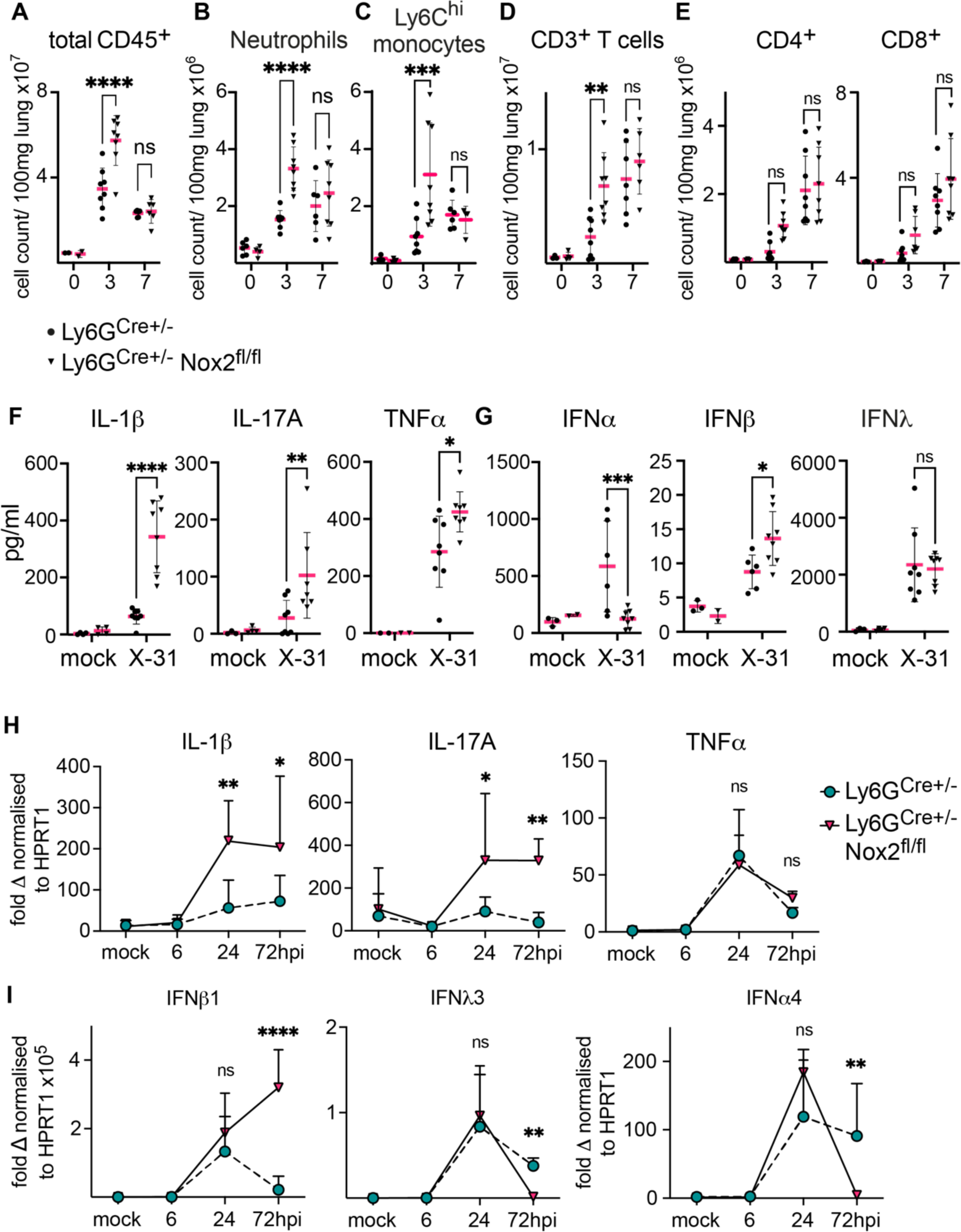
Increased activation of IL-1β /IL-17-axis in IAV-infected neutrophil-specific Nox2-deficient mice. *Ly6G-Cre*^+/−^ and *Ly6G-Cre*^+/−^ *Nox2^fl/fl^*mice were infected with 3×10^4^ TCID_50_ IAV strain X-31 or PBS (mock) in 30 µl intranasally. A – E) Number of Live CD45^+^ lymphocytes (A), Neutrophils gated as CD11b^hi^, Ly6G^tdTom^ (B), inflammatory Monocytes gated as CD11b^hi^, Ly6G^-^, Ly6C^hi^, MHCII^+^ (C), CD3e^+^ T cells (D), CD8^+^ and CD4^+^ αβ T cells (E) per 100 mg lung tissue 3 and 7 days post-infection (n = 3 - 4 mock, n = 4 - 8 X-31). F – G) Cytokines (F) and interferons (G) were quantified in bronchoalveolar lavage 3 days post-infection by ELISA (n = 2 - 3 mock, n = 8 X-31). H – I) Cytokine (H) and Interferon (I) mRNA expression in fold change from naïve normalised to HPRT1 in lung tissue was quantified at indicated time points by RT-qPCR (n = 3 mock, n = 6 X-31). Bars represent mean ± SEM of 2 – 3 independent experiments. p values are indicated, *<0.05, **<0.01, ***<0.001, ****<0.0001, ns not significant. p values were determined using two-way ANOVA and Sidak’s post-test (A - G) and Mann Whitney test (H - I).

Strikingly, overall chemokine and cytokine release was enhanced in the bronchoalveolar lavage (BAL) of IAV-infected *Ly6G-Cre*(+/−) *Nox2 flox* mice (Figure S3E and F). Specifically, chemokines and cytokines controlling the recruitment of neutrophils and monocytes as well as B and T cell proliferation and activation were elevated 3 days post-infection. While factors supporting resolution of inflammation were downregulated (e.g. Reg3G, EGF and FGF), indicating an overall dysregulated immune response in IAV-infected *Ly6G-Cre*(+/−) *Nox2 flox* mice. We confirmed increased presence of pro-inflammatory IL-1β, IL-17A and TNFα protein in BAL at day 3 (Figure 2F). Transcription of IL-1β and IL-17A mRNA increased at day 1 and remained elevated in *Ly6G-Cre*(+/−) *Nox2 flox* mice at day 3, while we observed no difference in TNFα transcription between both mouse strains (Figure 2H). Interestingly, while anti-viral interferon β1 (IFNβ1) correlated with increased IL-1β and IL-17A protein expression, IFNα4 and IFNλ3 expression quickly declined after initial up-regulation at day 1 in *Ly6G-Cre*(+/−) *Nox2 flox* mice (Figure 2G and I). In contrast, *Ly6G-Cre*(+/−) control mice showed only marginally increased IL-1β and IL-17A protein in BAL while IFNα4 and IFNλ3 protein and mRNA stayed elevated until day 3. Moreover, mRNA expression of adaptive type 2 IFNγ was reduced at day 3 post-infection in *Ly6G-Cre(+/−) Nox2 flox* mice (Figure S3G). Additionally, we observed elevated amounts of prostaglandin E2 (PGE2) protein in BAL of *Ly6G-Cre(+/−) Nox2 flox* compared to *Ly6G-Cre(-/-) Nox2 flox* control mice, a known inhibitor of type I interferon production and antiviral immunity induced by IL-1β (Figure S3H).^(28)^ In conclusion, neutrophil-specific Nox2-deficient mice shift their lung cytokine profile towards IL-1β/IL-17A production while the expression of anti-viral type I IFNα and type III IFNλ declined quickly, inversely correlating with virus burden.

### Enhanced γδ T cell-driven IL-17A response in Nox2-deficient mice

We performed intracellular cytokine staining to determine the cellular source of elevated proinflammatory cytokines. Mainly, neutrophils and inflammatory Ly6C^high^ monocytes were identified as IL-1β producers in *Ly6G-Cre*(+/−) *Nox2 flox* mice at day 3 post-infection (Figures 3A, B and S4A). Moreover macrophages, conventional dendritic cells and alveolar macrophages were found to produce IL-1β (Figures S4B, C and D). Notably, alveolar macrophages were also found to increase TNFα production in *Ly6G-Cre*(+/−) *Nox2 flox* mice (Figure S4E). Alveolar macrophage derived-TNFα has been reported previously to prime neutrophils for IL-1β production in a model of invasive pneumococcal disease^(29)^ and depletion of neutrophils through repeated injection of anti-Ly6G antibody in *Nox2-/-* mice confirmed neutrophils as main source of elevated IL-1β transcripts in lungs 3 days post-IAV infection (Figure S4F).

**Figure 3.**
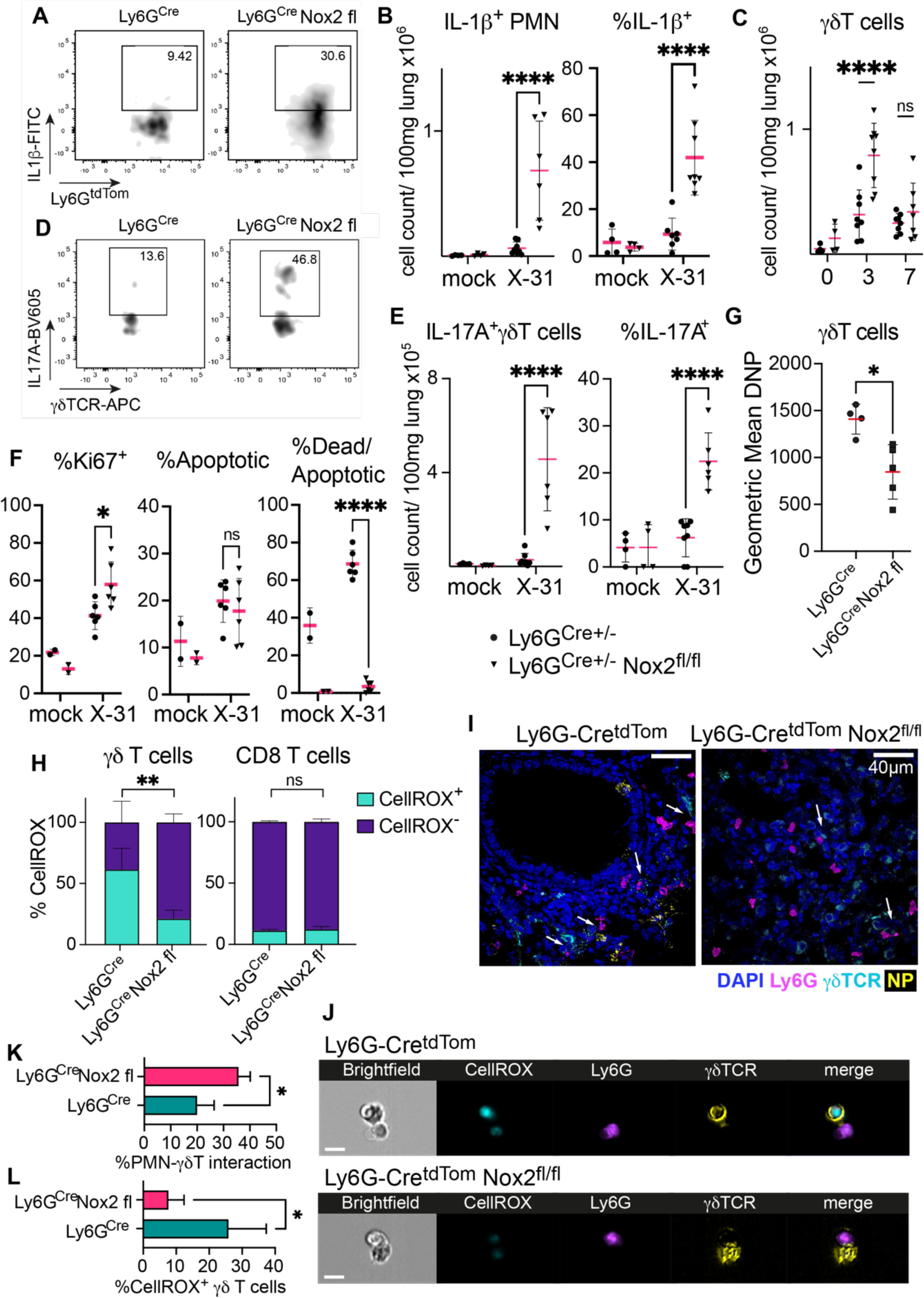
Enhanced γδ T cell-driven IL-17 response in IAV-infected Nox2-deficient animals. *Ly6G-Cre*^+/−^ and *Ly6G-Cre*^+/−^ *Nox2^fl/fl^*mice were infected with 3×10^4^ TCID_50_ IAV strain X-31 or PBS (mock) in 30 µl intranasally for indicated time points. A) Representative FACS plots of IL-1β^+^ neutrophils gated as Live CD45^+^, CD11b^+^ and Ly6G^tdTom^ at 3 days post-infection. B) Count per 100 mg lung tissue (left panel) and frequency (right panel) of IL-1β^+^ neutrophils at 3 days post-infection (n = 4 mock, n = 6 - 8 X-31). C) Number of γδ T cells gated as Live CD45^+^, CD3e^+^ and γδTCR^+^ per 100 mg lung tissue at 3 and 7 days post-infection (n = 4 mock, n = 8 X-31). D) Representative FACS plots of IL-17A^+^ γδ T cells at 3 days post-infection. E) Count per 100 mg lung tissue (left panel) and frequency (right panel) of IL-17A^+^ γδ T cells (n = 4 mock, n = 6 - 8 X-31). F) Frequency of Ki67^+^ (left panel), Apotracker^+^ (middle panel) and ZombieViolet^+^ Apotracker^+^ double-positive (right panel) γδ T cells at 3 days post-infection (n = 2 mock, n = 6 X-31). G) Protein oxidation in γδ T cells assessed by Dinitrophenylhydrazone (DNP)-staining at 3 days post-infection (n = 4 - 5). H) Frequency of CellROX^+^ γδ T (left panel, n = 6) and CD8 T cells (right panel, n = 3) at 3 days post-infection. I) Representative micrographs of lung sections stained for IAV nucleoprotein (NP, yellow), γδ T cells (γδTCR, cyan), neutrophils (Ly6G, magenta) and nuclei (DAPI, blue). Scale bar 40 µm. J - K) Representative imagestream graphs gated for Live CD45^+^ neutrophils (Ly6G^tdTom^, magenta), γδ T cells (γδTCR, yellow) and Oxidative stress (CellROX, cyan). Scale bar 10 µm (J). Quantitated in (K) Frequency of Live CD45^+^ γδ T cells in contact with neutrophils (n = 4). L) Frequency of Live CD45^+^ CellROX^+^ γδ T cells in contact with neutrophils (n = 4). Bars represent mean ± SEM of 2 – 3 independent experiments. p values are indicated, *<0.05, **<0.01, ****<0.0001, ns not significant. p values were determined using two-way ANOVA and Sidak’s post-test (B-F) and Mann Whitney test (G, H, K, L).

Neither CD4 nor CD8 αβ T cells were found to express significant amounts of IL-17A upon IAV infection in *Ly6G-Cre*(+/−) and *Ly6G-Cre*(+/−) *Nox2 flox* mice (Figure S4G and H). Since αβ T cells did not alone account for increased lung infiltration of CD3^+^ T cells at day 3 we further dissected the nature of lung-infiltrating T cells at this early time point. Strikingly, we observed a 2.5-fold elevated count of γδ T cells in *Ly6G-Cre*(+/−) *Nox2 flox* mice compared to *Ly6G-Cre*(+/−) mice which rapidly declined at day 7 (Figure 3C). Moreover, we identified γδ T cells as predominant source of elevated IL-17A at 3 days post-infection in *Ly6G-Cre*(+/−) *Nox2 flox* mice (Figure 3D and E). While the typical virus-induced type 1 IFNγ expression^(30)^ was slightly decreased in γδ T cells (Figure S4I) as well as in CD4 and CD8 αβ T cells (Figure S4G and H).

Surging γδ T cell counts correlated with increased cell proliferation (measured by Ki67 staining, left panel) and while the frequency of single-stained apoptotic γδ T cells (middle panel) was not affected, percentage of double-positive apoptotic dead cells (right panel) was significantly reduced in *Ly6G-Cre*(+/−) *Nox2 flox* mice (Figure 3F and Figure S5A and B).

Confirming the changes in γδ T cell proliferation and cytokine expression depend on neutrophil Nox2-mediated ROS production, we found that protein oxidation in γδ T cells was 0.5-fold reduced in *Ly6G-Cre*(+/−) *Nox2 flox* mice by analysing protein carbonylation (Figure 3G). Moreover, percentage of CellROX^+^ γδ T cells was 0.35-fold reduced in *Ly6G-Cre*(+/−) *Nox2 flox* mice (left panel) while CD8 (right panel), CD4 T cells and NK cells seemed unaffected by neutrophil ROS production with generally low percentage of CellROX^+^ cells (Figure 3H and S5C and D).

To substantiate a potential interaction between neutrophils and γδ T cells in IAV-infected lungs we verified both cell types co-localise near sites of infection (Figure 3I). Using ImageStream analysis we found Ly6G^tdTom^ neutrophils and γδ TCR^+^ T cells in direct cell contact (Figure 3J). Cell contact between both cells was increased in *Ly6G-Cre*(+/−) *Nox2 flox* mice (Figure 3K), while the percentage of CellROX^+^ γδ T cells in contact with neutrophils was significantly elevated in *Ly6G-Cre*(+/−) control mice (Figure 3L). In conclusion, neutrophils and γδ T cells directly interact in IAV-infected lungs, where Nox2-derived ROS induce oxidative damage and apoptosis in γδ T cells and moreover suppress the proliferation of γδ^17^ T cells.

### Nox2-deficient neutrophils induce γδ^17^ T cells

To elucidate the Nox2-dependent interaction between γδ T cells and neutrophils, we conducted *ex vivo* experiments using cells isolated from mouse tissues. We obtained neutrophils via negative selection from bone marrow (BM), achieving a purity of 85.23 ± 9.86 % neutrophils of live CD45^+^ cells (Figure S6A). Firstly, we confirmed assembly and activation of Nox2 in BM neutrophils upon infection with purified IAV by immunofluorescence staining (Figure 4A). We observed translocation of cytoplasmic Nox2 subunits p47phox (magenta) and p67phox (yellow) to the plasma membrane starting 30 minutes post-infection and significant colocalisation of both subunits measured by Pearson’s correlation 60 minutes post-infection (Figure 4B). We isolated γδ T cells from spleen and lymph nodes of wild-type mice (cell purity shown in Figure S6B) and co-cultured them with neutrophils from either wild-type or *Nox2-/-* mice. Direct cell contact was observed in co-cultures (Figure 4C), with a significant percentage of γδ T cells in contact with IAV-infected wild-type neutrophils staining CellROX^+^ (75 ± 36 %) as compared to 1.7 ± 0.26 % in contact with *Nox2-/-* neutrophils (Figure 4D and S7A).

**Figure 4.**
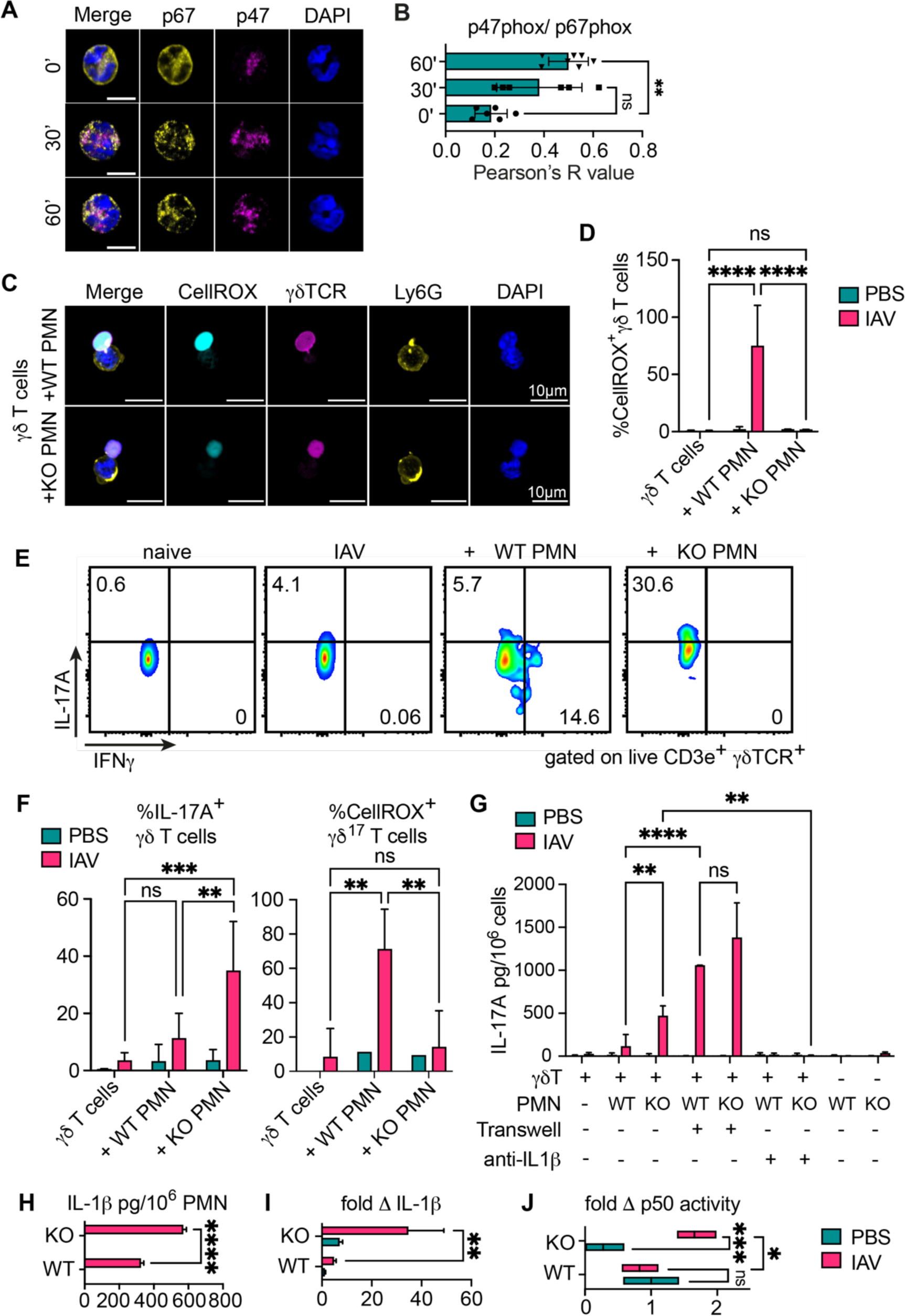
Nox2-deficient mouse neutrophils induce γδ^17^ T cells upon IAV infection. γδ T cells isolated from the spleen of naïve C57Bl/6J mice were co-cultured *ex vivo* with WT (C57Bl/6) or KO (Nox2-/-) neutrophils isolated from bone marrow at a ratio of 1:4 and were treated with PBS or infected with IAV strain PR8 at 0.2 MOI (multiplicity of infection). A) Representative micrographs of WT neutrophils at indicated time points (minutes) post-infection stained for p47-phox (magenta), p67-phox (yellow) and nuclei (DAPI, blue). Scale bar 5 µm. B) Pearson’s R values for colocalisation of p47- and p67-phox (n = 3 - 5). C) Representative micrographs of *ex vivo* co-cultures 2 hpi (hours post-infection) stained for Oxidative stress (CellROX, cyan), neutrophils (Ly6G, magenta), γδ T cells (γδTCR, yellow) and nuclei (DAPI, blue). Scale bar 10 µm. D) Frequency of CellROX^+^ γδ T cells 2 hpi cultured alone, in contact with WT or KO neutrophils (n = 3 - 4). E) Representative FACS plots of IL-17A^+^/IFNγ^+^ γδ T cells gated as Live CD45^+^, CD3e^+^, CD4^-^ and γδTCR^+^ 5 hours post-infection cultured alone, in contact with WT or KO neutrophils. F) Frequency of IL-17A^+^ γδ T cells (left panel) and CellROX^+^ γδ^17^ T cells (right panel) 5 hours post-infection cultured alone, in contact with WT or KO neutrophils (n = 4). G) IL-17A release assessed by ELISA 16 hours post-infection of γδ T cells, WT and KO neutrophils alone, γδ T cells co-cultured with WT or KO neutrophils, separated by transwell or pre-treated with anti-IL1β blocking antibody (n = 2). H) IL-1β release of WT and KO neutrophils assessed by ELISA 16 hours post-infection (n = 3). I) IL-1β mRNA expression of WT and KO neutrophils assessed by RT-qPCR 4 hours post-infection (n = 3). J) NFκB activity in lysates of WT and KO neutrophils 45 minutes post-infection (n = 4). Bars represent mean ± SEM of technical replicates representative of 2 - 4 independent experiments (B, G, H, I, J) or mean ± SEM of 3 - 4 independent experiments (D, F). p values are indicated, *<0.05, **<0.01, ***<0.001, ****<0.0001, ns not significant. p values were determined using Kruskal Wallis test with Dunn’s post-test (B), two-way ANOVA and Tukey’s post-test (D, F, G) and Sidak’s post-test (H, I, J).

Infection of isolated γδ T cells with IAV did not change their IL-17A or IFNγ expression (Figure 4E). However, co-culture with IAV-infected wild-type neutrophils slightly induced IFNγ production, while IAV-infected *Nox2-/-* neutrophils significantly shifted γδ T cells towards IL-17A production (Figure 4E), resulting in a 10-fold increase in IL-17A expression in co-cultured γδ T cells (Figure 4F, left panel). Moreover, Nox2-derived ROS production from wild-type neutrophils affected γδ^17^ T cells, as CellROX^+^ staining was 5-fold induced compared to co-culture with *Nox2-/-* neutrophils (Figure 4F, right panel). In addition, an increased percentage of γδ T cells expressed the IL-17 lineage-determining transcription factor RORγt in co-culture with *Nox2-/-* neutrophils (Figure S7B and C). In contrast, wild-type neutrophils tended to induce IFNγ production in γδ T cells (Figure S7D, left panel), and Nox2 presence did not significantly impact oxidative stress in γδ^IFNγ^ T cells (Figure S7D, right panel). These findings align with Mensurado et al., who previously showed that CD27^-^ γδ^17^ T cells are more susceptible to oxidative stress induced by neutrophil Nox2-mediated ROS production due to low transcription of antioxidant response genes.^(25)^

We further observed increased release of IL-17A in cell culture supernatant upon co-culturing with IAV-infected *Nox2-/-* neutrophils measured by ELISA (Figure 4G). Moreover, significantly elevated IL-17A mRNA expression in IAV-infected γδ T cells co-cultured with *Nox2-/-* neutrophils underlined a transcriptional regulation (Figure S7E). Of note, neutrophils alone did not produce significant levels of IL-17A. Using transwell inserts we demonstrated that Nox2-mediated suppression of IL-17A production was indeed dependent on direct cell contact. However, transwell experiments still showed higher induction of IL-17A release in co-culture with *Nox2-/-* neutrophils compared to wild-type neutrophils, indicating that *Nox2-/-* neutrophils signal to γδ T cells to proliferate and alter their cytokine profile. Since *in vivo* experiments showed elevated IL-1β production in lung neutrophils, we blocked IL-1β signalling in co-culture experiments by addition of neutralising antibody and completely abrogated IL-17A release from γδ T cells (Figure 4G).

We confirmed that *Nox2-/-* neutrophils release more IL-1β protein and mRNA upon IAV infection compared to wild-type (Figures 4H and I). Previous studies revealed a mechanism for Nox2-dependent regulation of neutrophil IL-1β production involving selective oxidation of key transcription factor NFκB,^(20)^ inhibiting its DNA-binding activity.^(18)^ We observed increased nuclear NFκB p50 subunit in lysates of IAV-infected *Nox2-/-* neutrophils (Figure S7F). To examine if this was sufficient to increase NFκB transcriptional activity, we employed a transcription factor-binding assay that measures the binding of active p50 to its consensus sequence. In contrast to infected wild-type neutrophils, p50 activity was significantly upregulated in *Nox2-/-* neutrophils (Figure 4J). In conclusion, we identified two Nox2-dependent mechanisms mediated by neutrophils that suppress early inflammatory responses to mild IAV infection. Firstly, a Nox2-mediated oxidative burst attenuates NFκB activity, reducing IL-1β production in neutrophils. Secondly, ROS suppress cytokine transcription in γδ^17^ T cells.

### Human neutrophils increase γδ^17^ T cells upon virus infection

A negative correlation between neutrophil and γδ T cell numbers has been noted in several human respiratory virus infections, suggesting an interaction between the two cell populations in humans as well. The ratio of immature neutrophils to γ(Vδ)2 T cells has been proposed to predict severe COVID-19.^(31)^ Further, in a human challenge study with RSV it was observed that high mucosal neutrophil numbers before exposure would predict symptomatic RSV disease with presymptomatic decline in mucosal IL-17A.^(32)^ We therefore explored whether our observations in mice could be extrapolated to human respiratory virus infections.

Firstly, we confirmed that virus infections induce ROS production via Nox2 activation in human neutrophils. We isolated neutrophils from the peripheral blood of healthy donors using centrifugation over a discontinuous density gradient and assessed purity by flow cytometry. On average 99.2 ± 0.5 % of Live CD45^+^ gated as CD11b^+^, CD14^-^, CD16^+^ neutrophils (Figure S8A). Nox2 assembly was observed by immunofluorescence staining for subunits p40phox (cyan), p47phox (magenta) and p67phox (yellow) 30 minutes post-infection with SARS-CoV-2 strain England 02 (Figure 5A, B and C) or stimulation with synthetic double-stranded RNA virus analogue poly(I:C) (polyinosinic-polycytidylic acid, data not shown). Transient colocalisation of subunits p40phox and p47phox with p67phox was confirmed by a significant rise in Pearson’s R value (Figure 5A and B). Additionally, we measured production of hydrogen peroxide with Oxyburst reagent in poly(I:C)-stimulated human neutrophils after 10 and 30 minutes (Figure 5D). We further explored the ROS burst induced by human neutrophils in response to virus infection with two chemiluminescent probes: luminol, which is cell permeable and detects both intracellular and extracellular ROS, and isoluminol, which detects only extracellular ROS. Poly(I:C) elicited a potent ROS burst within 20 – 60 minutes, peaking at 30 minutes, however very little extracellular ROS was measured (Figure S8C). The initial delay in ROS detection is consistent with ROS being consumed intracellularly.

**Figure 5.**
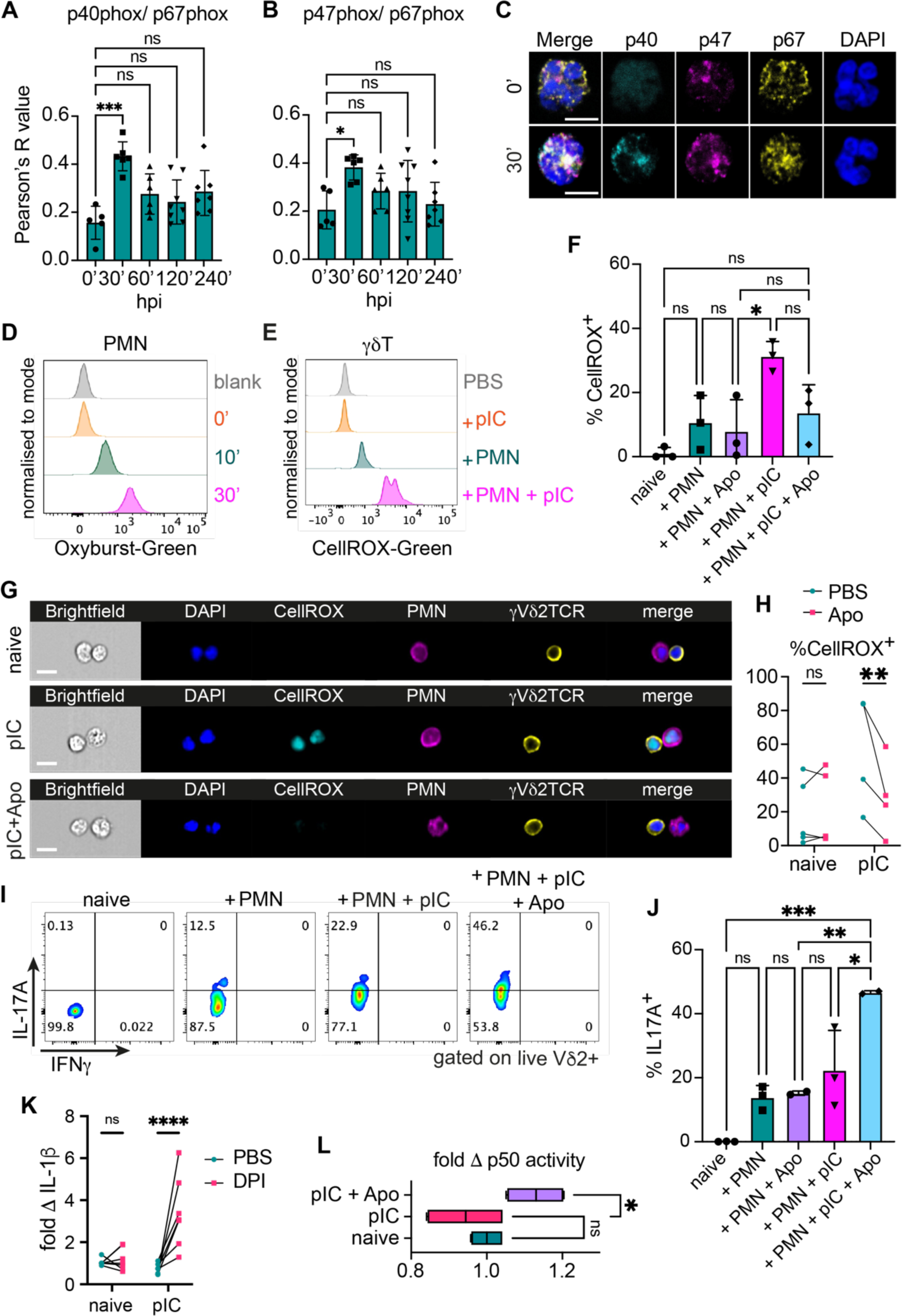
Human neutrophils enhance γδ^17^ T cells through Nox2-regulated IL-1β production during viral infection. γδ T cells and neutrophils isolated from peripheral blood of healthy volunteers were co-cultured *ex vivo*. A - B) Isolated human neutrophils were infected with SARS-CoV-2 at 0.2 MOI for indicated time points (minutes). Pearson’s R values for colocalisation of p40- and p67-phox (A) and p47- and p67-phox (B) (n = 5 – 8). C) Representative micrographs of isolated neutrophils 30 minutes post-infection stained for p40-phox (cyan), p47-phox (magenta), p67-phox (yellow) and nuclei (DAPI, blue). Scale bar 5 µm. D) Representative histogram plot of Oxyburst^+^ neutrophils stimulated for 0, 10 and 30 minutes with 1 µg/ml complexed poly(I:C) (pIC) gated as Live CD45^+^, CD11b^+^, CD16^+^ and CD14^-^. E - F) Representative histogram plot of CellROX^+^ γδ T cells cultured alone, in co-culture (ratio 1:10) with autologous unstimulated or pIC-stimulated neutrophils for 5 hours gated as Live CD45^+^, CD3^+^, CD4^-^ and γVδ2TCR^+^ cells (E). Quantitated in (F) as frequency of CellROX^+^ γδ T cells pre-treated 1 hour with PBS or 300 µM Apocynin (Apo) cultured alone, in co-culture with unstimulated or pIC-stimulated neutrophils for 5 hours (n = 3). G - H) Representative imagestream graphs of *ex vivo* co-cultures pre-treated 1 hour with PBS or 300 µM Apocynin (Apo), stimulated 5 hours with pIC and stained for Oxidative stress (CellROX, cyan), neutrophils (CD16, magenta), γδ T cells (γVδ2TCR, yellow) and nuclei (DAPI, blue). Scale bar 10 µm (G). Quantitated in (H) as frequency of CellROX^+^ γδ T cells in contact with neutrophils, values from same donor are indicated (n = 4 - 5). I) Representative FACS plots of IL-17A^+^/IFNγ^+^ γδ T cells pre-treated 1 hour with PBS or 300 µM Apocynin (Apo) cultured alone, in co-culture with unstimulated or pIC-stimulated neutrophils for 5 hours, gated as Live CD45^+^, CD3^+^, CD4^-^ and γVδ2TCR^+^ cells. J) Frequency of IL-17A^+^ γδ T cells pre-treated 1 hour with PBS or 300 µM Apocynin (Apo) cultured alone, in co-culture with unstimulated or pIC-stimulated neutrophils for 5 hours, gated as Live CD45^+^, CD3^+^, CD4^-^ and γVδ2TCR^+^ cells (n = 2 - 3). K) IL-1β release of human neutrophils pre-treated 1 hour with PBS or 10 µM Diphenyleneiodonium (DPI) and stimulated with 1 µg/ml pIC for 5 hours assessed by ELISA calculated as fold change from untreated. Values from same donor are indicated (n = 5). L) NFκB activity in human neutrophils pre-treated 1 hour with PBS or 300 µM Apocynin (Apo) and stimulated with 1 µg/ml pIC for 45 minutes calculated as fold change from untreated (n = 2). Bars represent mean ± SEM of technical replicates representative of 2 independent experiments (A, B) or mean ± SEM of 2 – 5 independent experiments (F, H, J, K, L). p values are indicated, *<0.05, **<0.01, ***<0.001, ****<0.0001, ns not significant. p values were determined using Kruskal Wallis test with Dunn’s post-test (A, B) two-way ANOVA with Tukey’s post-test (F, J), two-way ANOVA with Sidak’s post-test (H, K) or Kruskal-Wallis test with Dunn’s post-test (L).

Next, we isolated human γδ T cells from the peripheral blood of healthy donors, finding that on average 78 ± 11.9% of live CD45^+^ T cells were γδ2 T cells (Figure S8B). Poly(I:C) stimulation alone did not induce oxidative stress in γδ2 T cells as indicated by CellROX^+^ staining, however poly(I:C)-stimulated neutrophils significantly upregulated CellROX^+^ γδ2 T cells, which was reversed by pre-treating neutrophils with Nox2 inhibitor Apocynin (Figure 5E and F).

Using ImageStream analysis we found human neutrophils and γδ2 T cells in direct cell contact (Figure 5G). Quantitation of neutrophil-γδ2 T cell doublets revealed a significant Apocynin-dependent reduction of CellROX^+^ γδ2 T-neutrophil doublets (Figure 5H).

Flow cytometry analysis of neutrophil-γδ2 T cell co-cultures showed that neither poly(I:C) alone, unstimulated, Apocynin-treated nor poly(I:C)-stimulated neutrophils increased IL-17A or IFNγ production in γδ2 T cells (Figure 5I and J). However, blocking ROS production in poly(I:C)-stimulated neutrophils with Apocynin pre-treatment significantly increased IL-17A^+^ γδ2 T cells. This suggests that human neutrophils, like mouse neutrophils, signal to γδ T cells to upregulate IL-17A production, a process inhibited by ROS and restored upon Nox2 inhibition.

Nox2 inhibition in poly(I:C)-stimulated neutrophils increased IL-1β release (Figure 5K), and immunoblot analysis showed stabilisation of NFκB subunit p50 in lysates of poly(I:C)-stimulated human neutrophils pre-treated with Nox2 inhibitors Apocynin or Diphenyleneiodonium (Figure S8D). Moreover, NFκB p50 transcriptional activity was significantly upregulated upon Nox2 inhibition (Figure 5L).

Our results suggest that the interplay between neutrophils and γδ T cells in regulating inflammatory cytokine signalling, particularly IL-17A, is a conserved mechanism observed in both mice and humans. This interaction, modulated by Nox2-mediated ROS production in neutrophils, may play a pivotal role in the immune response to respiratory viral infections.

### Early Nox2-deficient neutrophil-derived IL-1β induces protective γδ^17^ T cell response in mice

Ultimately, we investigated the impact of neutrophil Nox2-mediated suppression of γδ^17^ T cells on the outcome of IAV infection in mice. We blocked IL-17 signalling by injecting an anti-IL17A neutralising antibody and compared virus burden 3 days post-infection to that of control IgG antibody-injected *Ly6G-Cre*(+/−) *Nox2 flox* mice. We observed a significant decrease in virus burden in IL-17A-depleted animals (Figure 6A), along with downregulation of IL-1β mRNA transcripts in the lungs (Figure 6B) and elevated levels of antiviral IFNα (Figure 6C), indicating a reinstated antiviral immune response.

**Figure 6.**
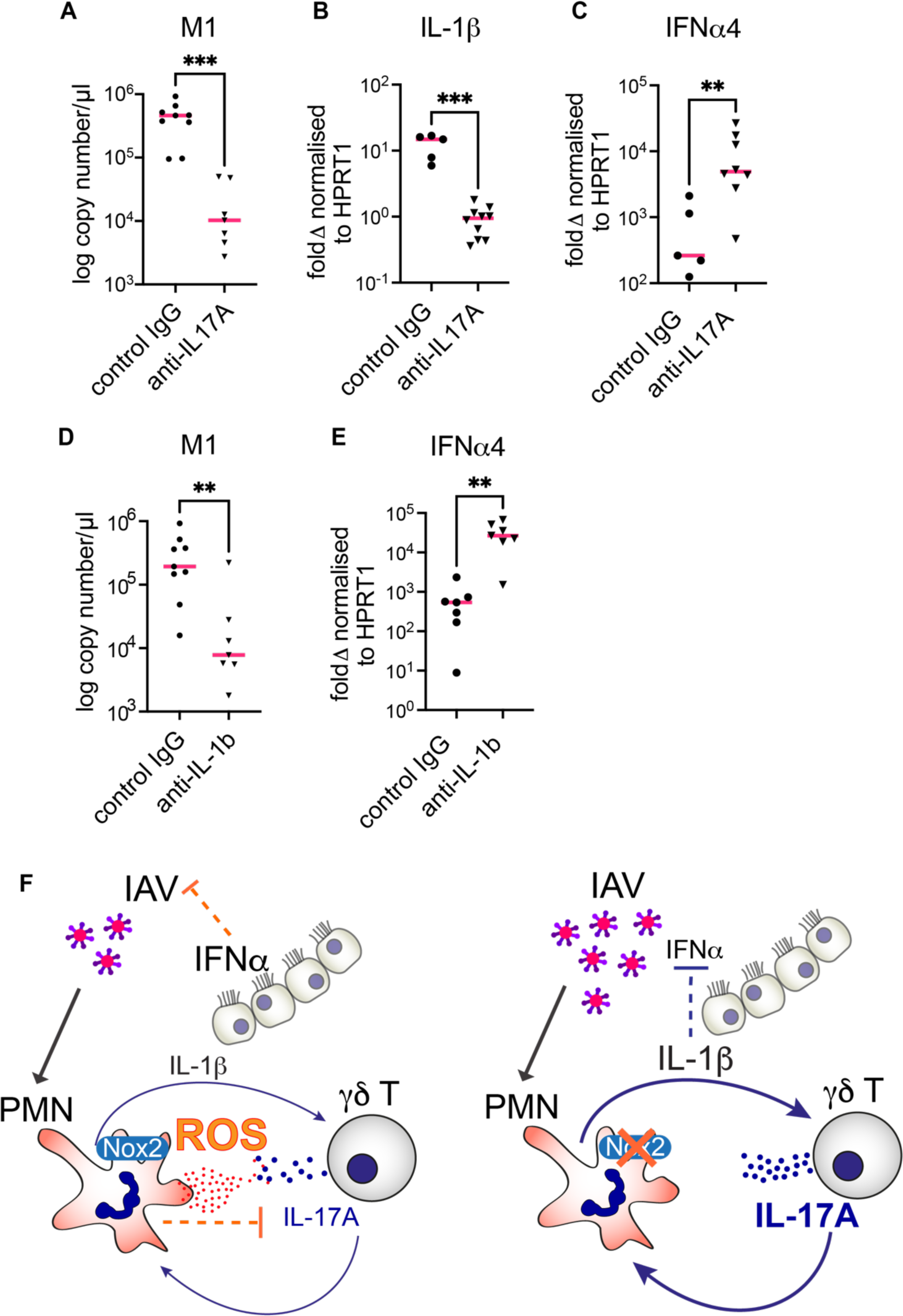
Early IL-1β production from Nox2-deficient neutrophils induces γδ^17^ T cell response and inhibits protective antiviral interferon response. *Ly6G-Cre*^+/−^ *Nox2^fl/fl^* mice were infected with 3×10^4^ TCID_50_ IAV strain X-31 or PBS (mock) in 30 µl intranasally for 3 days. A) Mice were treated with 100 µg anti-IL17A or control IgG antibody in 200 µl intraperitoneally at −1 day, −1 hour prior- and +1 day post-infection. Virus burden was assessed by RT-qPCR of influenza matrix protein M1 of whole lungs (n = 7 - 9). B - C) IL-1β (B) IFNα4 (C) mRNA expression in whole lung quantified by RT-qPCR (n = 5 - 10). E) Mice were treated with 100 µg anti-IL1β or control IgG antibody in 200 µl intraperitoneally 1 and 2 days post-infection. Virus burden was assessed by RT-qPCR of influenza matrix protein M1 of whole lungs (n = 7 - 9). F) IFNα4 mRNA expression in whole lung quantified by RT-qPCR (n = 7). G) Proposed model of neutrophil-γδ T cell interaction during early influenza A virus infection. Bars represent mean ± SEM of 2 - 3 independent experiments. p values are indicated, *<0.05, **<0.01, ***<0.001, ns not significant. p values were determined using Mann Whitney test.

Moreover, we reversed the viral burden by neutralising IL-1β signalling, which also correlated with increased IFNα production. In conclusion, neutrophil Nox2 plays a crucial role in controlling inflammation during virus infection and maintaining an effective antiviral immune response.

## Discussion

Our work explores the role of neutrophil Nox2 in regulating immune responses during respiratory viral infections, particularly focusing on the balance between inflammation and antiviral immunity. Neutrophil Nox2 is identified as the main source of ROS early after IAV infection. Early Nox2-derived ROS suppressed IL-1β signalling, which in turn regulated γδ T cell proliferation. In the absence of Nox2, unchecked IL-1β production by neutrophils led to an increase in IL-17-producing γδ T cells, resulting in an amplification of the IL-1β/IL-17 axis. This exacerbation interfered with the antiviral type I IFN response, resulting in diminished viral clearance and heightened lung inflammation. Additionally, we identified a self-perpetuating loop between IL-1β-producing neutrophils and IL-17-secreting γδ T cells, observed in both IAV-infected mice and human cells exposed to virus analogues.

Our study proposes a mechanism where neutrophil Nox2-derived ROS act as crucial modulators of cytokine signalling during viral infections. Prior research has demonstrated the role of the neutrophil ROS burst in tailoring inflammation based on the size of encountered microbes during bacterial and fungal infections.^(20)^ Small microbes are phagocytosed by neutrophils, triggering intracellular ROS production that suppresses IL-1β production through selective oxidation of transcription factor NFκB. This reduces IL-1β-mediated expression of neutrophil-recruiting chemokines KC and MIP-2 in mice, thus attenuating neutrophil recruitment and containing inflammation, as each neutrophil can eliminate numerous small pathogens. Conversely, large microbes trigger extracellular ROS production, enabling IL-1β production and subsequent neutrophil swarming to the infection site. This work has highlighted a critical role for the cellular localisation of ROS in regulating the recruitment of immune cells and their organisation in cooperative clusters. Defects in ROS production resulted in large neutrophil infiltrates in response to small microbes which would contribute to inflammatory disease.^(20)^

Our findings extend this knowledge to viral infections. We observed early assembly of the Nox2 complex involving p40phox, p47phox and p67phox subunits in endosomal compartments within 30 minutes of adding virus to neutrophil cultures. Moreover, we detected a strong ROS burst with the cell-permeable luminol probe, but not with cell-impermeable isoluminol, indicating rapid intracellular ROS production in response to virus recognition. In turn, this downregulated NFκB activity, which was reversed by Nox2 inhibitors. ROS-dependent regulation of NFκB binding to DNA has been demonstrated previously.^(18)^ However, we cannot rule out the involvement of redox-sensitive caspase-1 in regulating cleavage of pro-IL-1β into active IL-1β, which remains a potential area for further exploration.^(19)^

Concomitantly, we observed upregulated IL-1β production in neutrophils of *Nox2-/-* and *Ly6G-Cre(+/−) Nox2 fl/fl* mice post-IAV infection together with increased levels of neutrophil-recruiting chemokines and neutrophil swarming to sites of infection, which ultimately impaired virus clearance and caused lung damage. In turn, in the presence of Nox2, ROS suppress the production of IL-1β from neutrophils, thereby limiting the proliferation of IL-17-producing γδ T cells.

The interaction between neutrophils and γδ T cells emerged as a critical factor in modulating the immune response during viral infections. Our data show that neutrophil-derived IL-1β promotes the proliferation of IL-17-producing γδ T cells while Nox2-generated ROS act as a counterbalance suppressing this pathway. Moreover, direct cell contact between neutrophils and γδ T cells increased Nox2-dependent oxidative stress in γδ T cells, influencing their function and survival.

Our findings align with previous research showing that neutrophil-derived ROS suppress γδ^17^ T cell function. In fungal and bacterial lung infection models, neutrophilic NLRP3 inflammasome-dependent IL-1β secretion regulates γδ^17^ T cell responses,^(29)^ and neutrophil depletion increased IL-17A production by γδ T cells.^(33)^ Catalase addition to co-cultures of human blood-isolated neutrophils and γδ T cells to break down hydrogen peroxide, reversed γδ T cell suppression, unlike addition of PGE2 or arginase inhibitors.^(34)^ Interestingly, nitrogen-containing bisphosphonates (N-BP) are taken up by neutrophils and turned into isopentenyl pyrophosphate (IPP), which is known to activate γδ T cells. However, while N-BP-stimulated monocytes drive γδ T cell proliferation, neutrophils inhibited their activation acting as a regulatory checkpoint.^(24)^ The same authors suggested the release of hydrogen peroxide and serine proteases by neutrophils as inhibiting factors.^(24)^ Further studies have focussed on the role of neutrophil ROS production in γδ T cell inhibition. In various sterile and infectious models, including the tumour microenvironment, bacterial lung infections, and the autoimmune skin disease psoriasis, neutrophil-induced oxidative stress has been implicated in the suppression of γδ T cell activity.^(25,26,35)^ Anthony et al. discovered that excessive ROS production in superoxide dismutase 3 (SOD3)-deficient mice led to early neutrophil apoptosis, which in turn reduced IL-1β levels and limited γδ T cell proliferation.^(35)^ Mechanistically, Mensurado et al. demonstrated that low expression of the antioxidant glutathione in CD27^-^ γδ^17^ T cells rendered them particularly susceptible to neutrophil-derived ROS,^(25)^ explaining the shift from IFNγ to IL-17A production in our model upon IAV infection of *Ly6G-Cre(+/−) Nox2 fl/fl* mice.

We cannot exclude an additional role for neutrophil-released serine proteases such as neutrophil elastase in γδ T cell inhibition. In *in vitro* culture systems, neutrophil elastase has been shown to suppress the proliferation of blood-derived Vγ9δ2 T cells and to modulate their cytokine production.^(36,37)^ While we focused on the Nox2-ROS axis, other redox-regulated pathways may contribute to the overall immune response during viral infections. Future studies could investigate how ROS interact with various neutrophil-secreted factors to modulate γδ T cell activity and antiviral immunity.

Recently, Carissimo et al. showed an elevated neutrophil-to-Vδ2 T cell ratio correlating with severe COVID-19, suggesting that increased numbers of neutrophils could cause γδ T cell deficiency in patients contributing to the dysregulated immune response.^(31)^ A human challenge study with RSV found that neutrophilic inflammation in the nasal mucosa predisposes to RSV infection, which was linked to a decline of mucosal IL-17.^(32)^ Interestingly, in our model increased IL-17 production in the absence of neutrophil Nox2 amplified viral burden and inflammation. This highlights the differing effects of IL-17 in the upper - especially nasal mucosa - versus lower respiratory tract, raising the question of when IL-17 becomes detrimental during IAV infection by impairing lung barrier function. The interplay between neutrophils and γδ T cells is crucial in determining the balance between effective antiviral responses and excessive inflammation, underscoring the potential for targeted therapeutic interventions to modulate these interactions.

Moreover, proinflammatory IL-1β/IL-17 signalling can downregulate the antiviral type I IFN response. Previous studies have shown that IL-1β-driven upregulation of PGE2 inhibits type I IFN production during IAV infection in mice. When PGE2 receptors on macrophages are activated, they trigger the PI3K-Akt pathway, which prevents the phosphorylation of IRF3, a crucial transcription factor for type I IFN production. This suppression leads to increased viral replication and a weakened antiviral immune response.^(28,38)^ Elevated PGE2 levels, correlating with increased IL-1β signalling and viral load, were also observed in IAV-infected *Ly6G-Cre(+/−) Nox2 fl/fl* mice.

Our study highlights the dual role of neutrophil-derived ROS - particularly Nox2-generated - in modulating immune responses. Traditionally viewed as damaging agents, ROS are demonstrated to play a critical role in immune regulation by suppressing inflammatory cytokine signalling (IL-1β/IL-17), modulating the activity and proliferation of γδ T cells, and balancing the antiviral interferon response to ensure efficient viral clearance while minimizing tissue damage.

This dual function positions Nox2-derived ROS as potential therapeutic targets, where modulating ROS levels could enhance antiviral immunity while controlling excessive inflammation. Targeting ROS production for therapeutic purposes requires a nuanced understanding of its localisation and context within the immune response. Neutrophil-specific Nox2-derived ROS have distinct roles compared to ROS produced by other cells and the localisation of ROS production within neutrophils is crucial for its regulatory effects on IL-1β and IL-17 signalling. This cell-specific role highlights the importance of developing therapies that focus on targeted regulation of ROS, rather than broad-spectrum approaches.

Broad-spectrum antioxidants may lack efficacy because they fail to discriminate between beneficial and harmful ROS. Therapeutic strategies should aim to target specific sources of ROS, such as Nox2 in airway epithelial versus neutrophils, to modulate the immune response effectively without broadly suppressing ROS needed for pathogen clearance and immune regulation. Spatial control of ROS production would enable precise modulation of specific immune pathways. For example, restricting antioxidant to the airway epithelium at virus entry could alleviate damage to the lung epithelium caused by oxidative stress without impairing the overall oxidative burst required for antiviral defence and control of excessive inflammation.

Our results are consistent with observations made in Chronic Granulomatous Disease (CGD) patients, who have genetic defects in Nox2 expression. CGD patients experience recurring bacterial and fungal infections and are also prone to develop chronic inflammation and autoimmune symptoms.^(39,40)^ Moreover, an expansion of Th17 cells has been particularly noted in CGD patients, further contributing to immune dysregulation.^(41,42)^

Further supporting the importance of cell-specific ROS regulation, studies using full Nox2 KO mice showed a modest improvement in the resolution of IAV infection, but with increased inflammation.^(15,16)^ These findings suggest that the absence of Nox2 enhances viral clearance at the expense of heightened inflammatory responses due to the loss of ROS-mediated regulation across all cell types. In contrast, our study using neutrophil-specific Nox2 KO mice reveals that neutrophil-derived ROS specifically regulate IL-1β and IL-17 signalling, playing a critical role in maintaining a balanced inflammatory response while preserving the antiviral functions of other immune cells. This comparison underscores the importance of cell-specific studies in understanding the nuanced roles of ROS in immune regulation.

A limitation of our study is the complexity of translating findings from murine models to human disease. Although we observed similar mechanisms in human cells exposed to viral analogues, clinical studies are needed to validate the therapeutic potential of targeting Nox2-derived ROS in humans. Furthermore, the context-dependent role of IL-17, which can be both protective and detrimental depending on the site of inflammation (upper versus lower respiratory tract), warrants further investigation to delineate its precise role in respiratory viral infections.

Our findings have significant implications for chronic respiratory diseases like asthma and Chronic Obstructive Pulmonary Disease (COPD), where oxidative stress and inflammation play key roles. The dual role of ROS suggests that modulating ROS levels through targeted interventions could help balance inflammation and enhance antiviral immunity, potentially reducing exacerbations in asthma and COPD patients. Recognizing the beneficial role of ROS in immune regulation supports the concept of “good oxidants”,^(43)^ where controlled ROS production is essential for maintaining immune homeostasis. The limited efficacy of broad-spectrum antioxidants in respiratory diseases can be explained by the need for a balanced ROS environment. Over-suppression of ROS might hinder essential immune functions, highlighting the need for targeted antioxidant therapies.

## Acknowledgements

The authors thank John McCauley for kindly providing Influenza A virus preparation, Rui Pedro Ribeiro Galao for SARS-Co-V2 infections. We thank Venizelos Papayannopoulos, Matthias Gunzer, Leo Caslin and Ajay Shah for kindly providing mouse models. We kindly thank the staff of the Nikon Imaging Centre, the BRC flow facility and Biological Service Unit NHH at King’s College London for technical advise. This Work was supported by the Wellcome Trust (Sir Henry Dale fellowship 216373/Z/19/Z).

## Author contributions

Conceptualisation, A.W.; Methodology, A.W.; Formal Analysis, A.S. and A.W.; Investigation, A. S., A. P., F.S., A. G., Z.A. and A. W.; Writing – Original Draft, A. W.; Writing – Review & Editing, A. W.; Visualisation, A.W.; Funding Acquisition, A.W.

## Declaration of generative AI and AI-assisted technologies in the writing process

During the preparation of this work the author used AI language model ChatGPT 4 from OpenAI to edit this document for brevity and clarity. After using this tool/service, the author reviewed and edited the content as needed and takes full responsibility for the content of the publication.

