## Supplementary material for "Neutrophil NADPH oxidase breaks the inflammatory IL-1β/IL-17A circuit to enhance pathogen clearance during respiratory virus infections": Methods

### STAR Methods

#### Key resources table

| Reagent or Resource | Source | Identifier (Cat.no., RRID) |
|---|---|---|
| **Antibodies** | | |
| Anti-goat IgG Alexa Fluor 488 | Invitrogen | Cat# A11055, RRID:AB2534102 |
| Anti-hamster IgG Alexa Fluor 647 | Invitrogen | Cat# A-21451, RRID:AB2535868 |
| Anti-human CD3-PE-Cy5 | Biolegend | Cat# 317355, RRID:AB2904343 |
| Anti-human CD4 BV650 | Biolegend | Cat# 300535, RRID:AB2632791 |
| Anti-human CD8a BV785 | Biolegend | Cat# 301045, RRID:AB2563264 |
| Anti-human CD11b PE-Cy5 | Biolegend | Cat# 301308, RRID:AB314160 |
| Anti-human CD14 BV786 | Biolegend | Cat# 301840, RRID:AB2563425 |
| Anti-human CD16 PE | Biolegend | Cat# 302056, RRID:AB314207 |
| Anti-human CD45 APC-Cy7 | Biolegend | Cat# 304014, RRID:AB314402 |
| Anti-human IL-1β Alexa Fluor 647 | Biolegend | Cat# 511707, RRID:AB2124352 |
| Anti-human IL-17A BV605 | Biolegend | Cat# 512326, RRID:AB2563887 |
| Anti-human IFNγ BV711 | BD Bioscience | Cat# 564039, RRID:AB2738557 |
| Anti-human/mouse Myeloperoxidase | R&D | Cat# AF3667, RRID:AB2250866 |
| Anti-human TCR Vδ2 APC | Biolegend | Cat# 331418, RRID:AB2687323 |
| Anti-Influenza A nucleoprotein (NP) | Invitrogen | Cat# PA5-32242, RRID:AB2549715 |
| Anti-mouse CD3e PE-Cy5 | Biolegend | Cat# 100310, RRID:AB312674 |
| Anti-mouse CD4 BV650 | Biolegend | Cat# 100546, RRID:AB2562098 |
| Anti-mouse CD8a PE-Cy7 | Biolegend | Cat# 100722, RRID:AB312746 |
| Anti-mouse CD11b APC | Biolegend | Cat# 101212, RRID:AB312795 |
| Anti-mouse CD11c PE-Cy7 | Biolegend | Cat# 117317, RRID:AB493569 |
| Anti-mouse CD19 BV650 | Biolegend | Cat# 115541, RRID:AB11204087 |
| Anti-mouse/human CD27 PE/Dazzle 594 | Biolegend | Cat# 124228, RRID:AB2565794 |
| Anti-mouse CD45 APC-Cy7 | Biolegend | Cat# 103116, RRID:AB312981 |
| Anti-mouse CD62L PE/Dazzle 594 | Biolegend | Cat# 104447, RRID:AB2566162 |
| Anti-mouse CD103 BV711™ | Biolegend | Cat# 121435, RRID:AB2686970 |
| Anti-mouse CD170 BV421 | Biolegend | Cat# 155509, RRID:AB2810421 |
| Anti-mouse/human Ki-67 BV421 | Biolegend | Cat# 151208, RRID:AB2629748 |
| Anti-mouse IgG Alexa Fluor 568 | Fisher Scientific | Cat# A10037, RRID:AB11180865 |
| Anti-mouse I-A/I-E PE-Cy5 | Biolegend | Cat# 107611, RRID:AB313326 |
| Anti-mouse IL-1β DyLight® 488 | Universal Biologicals Ltd | Cat# I-960, RRID:AB28310 |
| Anti-mouse IL-17A BV605™ | Biolegend | Cat# 506927, RRID:AB11126144 |
| Anti-mouse IFN-γ BV711 | Biolegend | Cat# 505836, RRID:AB2650928 |
| Anti-mouse Ly-6G PE | Biolegend | Cat# 127608, RRID:AB1186099 |

| | | |
|---|---|---|
| Anti-mouse NK1.1 Pacific Blue | Biolegend | Cat# 108721, RRID:AB2234352 |
| Anti-mouse Rorγt APC | R&D | Cat# IC9125A-025 |
| Anti-mouse TCR γ/δ APC | Biolegend | Cat# 118116, RRID:AB1731813 |
| Anti-mouse TCR Vγ 1.1 APC | Biolegend | Cat# 141107, RRID:AB1089780 |
| Anti-NCF2, p67-phox | mybiosource | Cat# MBS8245577 |
| Anti-NCF4, p40-phox | mybiosource | Cat# MBS127688 |
| Anti-NFKB p105/ p50 [E381] | Abcam | Cat# ab32360, RRID:AB776748 |
| Anti-p47-phox [D-10] | Santa Cruz | Cat# sc-17845, RRID:AB627986 |
| Anti-rabbit IgG, Alexa 568 | Invitrogen | Cat# A10042, RRID:AB2534017 |
| Human TruStain FcX | Biolegend | Cat# 422302, RRID:AB2818986 |
| InVivoPlus Anti-mouse IL-1β | BioXcell | Cat# BE0246, RRID: AB2687727 |
| InVivoPlus Anti-mouse IL-17A | BioXcell | Cat# BE0173, RRID:AB10950102 |
| InVivoMab Anti-mouse TCR γ/δ | BioXcell | Cat# BE0070, RRID:AB1107751 |
| InVivoPlus Anti-mouse Ly6G | BioXcell | Cat# BE0075, RRID:AB1107721 |
| InVivoPlus polyclonal Armenian hamster IgG | BioXcell | Cat# BE0091, RRID:AB1107773 |
| InVivoPlus mouse IgG1 isotype control | BioXcell | Cat# BE0083, RRID:AB1107784 |
| InVivoPlus rat IgG2a isotype control | BioXcell | Cat# BE0089, RRID:AB1107769 |
| TruStain FcX (Anti-mouse CD16/32) | Biolegend | Cat# 101320, RRID:AB1574975 |
| **Virus strains** | | |
| Influenza A/ HKx31 (X-31) H3N2 | John McCauley (Francis Crick Institute) | N/A |
| Influenza A/ Puerto Rico/ 8/ 34 (PR8) H1N1 | John McCauley (Francis Crick Institute) | N/A |
| SARS-CoV2 England 02/2020/407073 | Rui Pedro Galao (King's College London) | N/A |
| **Chemicals, reagents, peptides and recombinant proteins** | | |
| Ammonium-chloride-potassium (ACK) buffer | Gibco | Cat# A104920 |
| Animal-Free Blocker and Diluent | 2B Scientific | Cat# SP-5035-100 |
| Apocynin | Merck | Cat# A10809 |
| Apotracker™ Green | Biolegend | Cat# 427402 |
| LIVE/DEAD™ Fixable Aqua Dead Cell Stain Kit | Invitrogen | Cat# L34957 |
| Bicinchoninic acid (BCA) protein assay kit | Pierce | Cat# 23227 |
| Bovine serum albumin (BSA) | Thermo Fisher | Cat# BP1600-100 |
| Brefeldin | Biolegend | Cat# 420601 |
| CBA Mouse/Rat Soluble Protein Master Buffer Kit | BD Bioscience | Cat# 558266 |
| Cell activation cocktail | Biolegend | Cat# 423302 |

| | | |
|---|---|---|
| CellROX™ Green Reagent, for oxidative stress detection | Invitrogen | Cat# C10444 |
| Complete Mini protease inhibitor cocktail | Roche | Cat# 1836153001 |
| Complexed Polyinosinic:polycytidylic acid (Poly-IC) | Invivo Gen | Cat# tlrl-picwlv |
| Fixation buffer | Biolegend | Cat# 420801 |
| Cytoperm/Cytofix buffer | BD Bioscience | Cat# 554714 |
| Dihydroethidium Reagent (DHE) | Invitrogen | Cat# D11347 |
| Diphenyleneiodonium chloride (DPI) | Cambridge Bioscience | Cat# 81050 |
| Dulbecco's modified Eagle medium (DMEM) | Gibco | Cat# 10564011 |
| EasySep human γ/δ T cell isolation kit | Stemcell Technologies | Cat# 19255 |
| EasySep Mouse Neutrophil enrichment kit | Stemcell Technologies | Cat# 19762 |
| EasySep Release Mouse APC positive selection kit | Stemcell Technologies | Cat# 100-0033 |
| Enhanced Chemiluminescence (ECL) substrate | Pierce | Cat# 35055 |
| Epoxomicin | Generon | Cat# A2606 |
| Fetal bovine serum (FBS) | Cytiva Hyclone | Cat# 12694237 |
| Fixation/Permeabilisation kit | BD Bioscience | Cat# 554714 |
| GlutaMAX™ Supplement | Gibco | Cat# 35050061 |
| Hank's Balanced Salt solution (HBSS) with $Ca^{2+}$ and $Mg^{2+}$ | Gibco | Cat# 14025092 |
| Hank's Balanced Salt solution (HBSS) without $Ca^{2+}$ and $Mg^{2+}$ | Gibco | Cat# 14170112 |
| HEPES 4-(2-hydroxyethyl)-1-piperazineethanesulfonic acid | Gibco | Cat# 15630056 |
| Histopaque 1119 | Merck | Cat# 11191-100ML |
| Human IL-1β Uncoated ELISA kit | Invitrogen | Cat# 88-7261-88 |
| Horse radish peroxidase HRP | Merck | Cat# 1162160001 |
| Image-iT™ fixative solution | Invitrogen | Cat# FB002 |
| Iscove's Modified Dulbecco's Medium (IMDM) | Gibco | Cat# 12440053 |
| Isoluminol | Sigma | Cat# A8264-1G |
| Liberase TL | Roche | Cat# 5401020001 |
| Luminol | Sigma | Cat# 123072-2.5G |
| Minimum essential medium (MEM) | Gibco | Cat# 41090028 |
| Monensin | Biolegend | Cat# 420701 |
| Mouse IFNα ELISA kit | Invitrogen | Cat# BMS6027TWO |

| | | |
|---|---|---|
| Mouse IFNβ ELISA kit | PBL | Cat# 42410-1 |
| Mouse IFNγ CBA Flex set A4 | BD Bioscience | Cat# 558296 |
| Mouse IFNγ ELISA kit | Invitrogen | Cat# 88-7314-22 |
| Mouse IFNλ ELISA kit | Biotechne | Cat# DY1789B-05 |
| Mouse IL-1β CBA Flex set E5 | BD Bioscience | Cat# 560232 |
| Mouse IL-1β Uncoated ELISA kit | Invitrogen | Cat# 88-7013-22 |
| Mouse IL-17A CBA Flex set C5 | BD Bioscience | Cat# 560283 |
| Mouse IL-17A Uncoated ELISA Kit | Invitrogen | Cat# 88-7371-22 |
| Mouse IL-17F CBA Flex set D6 | BD Bioscience | Cat# 562174 |
| Mouse IL-6 CBA Flex Set B4 | BD Bioscience | Cat# 558301 |
| Mouse KC CBA Flex set A9 | BD Bioscience | Cat# 558340 |
| Mouse MCP-1 CBA Flex set B7 | BD Bioscience | Cat# 558342 |
| Mouse PGE2 ELISA kit | LSBio | Cat# LS-F32354-LSP |
| Mouse TNFα CBA Flex set C8 | BD Bioscience | Cat# 558299 |
| Mouse TNFα ELISA kit | Stemcell Technologies | Cat# 2030 |
| N-Ethylmaleimide | Cambridge Bioscience | Cat# HY-D0843 |
| Nuclear Extraction kit | Cambridge Bioscience | Cat# 10009277 |
| NuPAGE 4-12% Bis Tris protein gel | Invitrogen | Cat# 10247002 |
| Optimal cutting temperature (OCT) medium | VWR | Cat# 361603E |
| Oxyburst™ Green H2DCFDA | Invitrogen | Cat# D-2935 |
| Paraformaldehyde 4% (PFA) | VWR | Cat# 9713 |
| Phosphate-buffered saline (PBS) | Gibco | Cat# 10010023 |
| PCRmax Human Influenza Type A M1 Kit | PCR Max | Cat# PKIT10035 |
| Percoll | GE Healthcare | Cat# 17544502 |
| Poly-D-lysine BioCoat | Corning | Cat# 354210 |
| ProLong™ Diamond Antifade Mountant | Invitrogen | Cat# P36970 |
| Proteome profiler mouse XL cytokine array | R&D | Cat# ARY028 |
| Recombinant human TNFα | Abcam | Cat# ab259410 |
| RNAlater™ Stabilisation solution | Invitrogen | Cat# AM7020 |
| RNA-to-CT™ TaqMan™ RNA-to-$C_T$™ *1-Step* Kit | Applied Biosystems | Cat# 10158014 |
| RNeasy kit | Qiagen | Cat# 74104 |
| TaqMan™ Universal Master Mix II, no UNG | Applied Biosystems | Cat# 4440047 |
| TransAM® NFκB p50 assay kit | Active Motif | Cat# 41096 |
| Transcriptor Frist Strand cDNA synthesis kit | Roche | Cat# 4379012001 |

| | | |
|---|---|---|
| TPCK-treated Trypsin | 2BScientific | Cat# 21560034-1 |
| TRI Reagent™ Solution | Invitrogen | Cat# AM9738 |
| Zombie Violet | Biolegend | Cat# 423113 |
| 4',6-diamidino-2-phenylindole (DAPI) | Insight Biotechnologies | Cat# AR1176 |
| **Experimental models: Cell lines** | | |
| Madin-Darby canine kidney (MDCK) cell line (NBL-2) | ATCC | RRID: CVCL_0422 |
| **Experimental models: Organism** | | |
| Mouse: C57BL/6-Ly6G(tm2621(Cre-tdTomato)Arte; Ai14 | Matthias Gunzer (University Duisburg-Essen) | Hasenberg et al. Nat Methods. 2015 doi: 10.1038/nmeth.3322. Epub 2015 Mar 16. PMID: 25775045. |
| Mouse: Nox2 fl/fl | Ajay Shah (King's College London) | Sag et al. Circulation 2017 doi: 10.1161/CIRCULATIONAHA.116.023877. PMID:28298457. |
| Mouse: Nox2 KO, B6.129S-Cybbtm1Din/J | Mary Dinauer (University St. Louis) | Pollock et al. Nat Genet 1995 https://doi.org/10.1038/ng0295-202 IMSR_JAX:002365 |
| Mouse: C57BL/6J | Jackson Laboratory | IMSR_JAX:000664 |
| **Oligonucleotides** | | |
| Human HPRT1 Hs02800695_m1 | Applied Biosystems | Cat# 4331182 |
| Human IL-1β Hs01555410_m1 | Applied Biosystems | Cat# 4331182 |
| Mouse B2M Mm00437762_m1 | Applied Biosystems | Cat# 4331182 |
| Mouse HPRT1 Mm03024075_m1 | Applied Biosystems | Cat# 4331182 |
| Mouse IFIT3 Mm01704846_s1 | Applied Biosystems | Cat# 4331182 |
| Mouse IFNα4 Mm00833969_s1 | Applied Biosystems | Cat# 4331182 |
| Mouse IFNβ1 Mm00439552_s1 | Applied Biosystems | Cat# 4331182 |
| Mouse IFNγ Mm01168134_m1 | Applied Biosystems | Cat# 4331182 |
| Mouse IFNλ3 Mm00663660_g1 | Applied Biosystems | Cat# 4331182 |
| Mouse IL-1β Mm00434228_m1 | Applied Biosystems | Cat# 4331182 |
| Mouse IL-17A Mm00439618_m1 | Applied Biosystems | Cat# 4331182 |
| Mouse ISG15 Mm01705338_s1 | Applied Biosystems | Cat# 4331182 |
| Mouse OAS Mm00836412_m1 | Applied Biosystems | Cat# 4448892 |
| Mouse Rsad2 Mm00491265_m1 | Applied Biosystems | Cat# 4331182 |
| Mouse Stat1 Mm01257286_m1 | Applied Biosystems | Cat# 4331182 |
| Mouse TNFα Mm00443258_m1 | Applied Biosystems | Cat# 4331182 |
| **Software and Algorithms** | | |
| GraphPad Prism v10 | GraphPad by Dotmatics | https://www.graphpad.com/ |
| FlowJo Version v10.9 | Becton, Dickinson and Company | https://www.flowjo.com/ |

| | | |
|---|---|---|
| Amnis Imagestream IDEAS software | Amnis Corporation | https://www.luminexcorp.com/imagestreamx-mk-ii/#software |
| Amnis Imagestream INSPIRE software | Amnis Corporation | https://www.luminexcorp.com/imagestreamx-mk-ii/#software |
| ImageJ 1.54f | Schneider et al., 2012/ NIH | https://imagej.org |
| Jupyterlab v3.6.7 | Project Jupyter | Jupyter.org |
| Python 3.9.19 | Python software foundation | |
| Colocalisation script | Annika Warnatsch | Supplemental file, GitHub Repository AnniWarn/Colocalisation |

#### Resource availability

##### Lead contact

##### Material availability

All newly generated materials associated with the paper are available upon request from the lead contact.

##### Data availability

- Code for colocalisation analysis can be assessed on GitHub Repository AnniWarn/Colocalisation
- Any additional information required to reanalyse the data reported is available from the lead contact upon request.

#### Experimental model and subject details

##### Mice

All mice strains used were derived from the C57Bl/6J background. Wild-type (WT) mice were purchased from The Jackson Laboratory. Nox2-/- mice were kindly provided by Venizelos Papayannopoulos (Francis Crick Institute) and described previously (Pollock et al. 1995), Nox2 fl/fl mice by Ajay Shah (King's College London)(Sag et al. 2017) and Ly6G-Cre/tdTomato mice by Matthias Gunzer (University Duisburg-Essen)(Hasenberg et al. 2015). Ly6G-Cre and Nox2 fl/fl mice were crossed to generate Ly6G-Cre(+/-) Nox2 fl/fl and Ly6G-Cre(-/-) Nox2 fl/fl mice. Colonies were maintained at the Biological Services Unit, King's College London, UK. All mice were housed in specific-pathogen free (SPF) conditions in full compliance with FELASA recommendations. Mice were housed at 22°C with a 12-hour light-dark cycle and given standard food and water ad libitum. Animals were 8-10 weeks old and age-matched within 1-2 weeks for experiments. Littermates of both sexes were randomly assigned to experimental groups. All procedures were approved by the Home Office under the Animals Scientific Procedures Act 1986 (ASPA).

**Human volunteers**

Human peripheral blood was obtained from healthy adult donors with informed consent. The study was approved by the ethical committee (REC14/LO/1699). The inclusion criteria of the donor were individuals 18 years of age or older and had no history of respiratory disease.

**Viruses**

Influenza A virus strains X-31 and PR8 were kindly provided by John McCauley (Francis Crick Institute). SARS-CoV2 England 02 strain was kindly provided by Rui Pedro Ribeiro Galao (King's College London).

#### Method details

**Virus purification and titration.** Influenza A virus strains were grown in chicken eggs, filtered, and titrated. $TCID_{50}$ was determined using Madin-Darby Canine Kidney (MDCK) cells. MDCK cells were seeded in 96-well plates in Dulbecco's Modified Eagle Medium (DMEM) supplemented with 10% fetal calf serum (FCS), 100 U/mL penicillin and 100 µg/mL streptomycin. At 80-90% confluency, plates were washed with phosphate-buffered saline (PBS), and 8-fold serially diluted virus was added in DMEM supplemented with 0.1% FCS, 0.3% bovine serum albumin (BSA), 100 U/mL penicillin and 100 µg/mL streptomycin, 200 µM GlutaMAX™, 20 mM HEPES and 1 µg/ml L-1-tosylamido-2-phenylethyl chloromethyl ketone (TPCK)-treated trypsin. Cells were fixed in 4% paraformaldehyde (PFA) after 5 days and stained with crystal violet. Plates were analysed for cytopathic effect and $TCID_{50}$ calculated using Reed and Muench methodology (Reed and Muench, 1938).

Influenza A virus was purified as described previously (Hutchinson et al., 2012). Briefly, MDCK cells grown in T175 flasks were inoculated at Multiplicity of Infection (MOI) 0.001 in 6 ml serum-free Minimum Essential Medium (MEM). Cells were incubated for 1 hour and 25 ml serum-free MEM was added per flask and incubated for 3 days. Supernatant was filtered and layered above 5 ml of 30% sucrose. Ultracentrifugation was carried out at 25,000 rpm (acceleration 9, deceleration 0) for 90 minutes at 4°C. Supernatant was removed and virus resuspended in 3 ml DMEM.

**Infection of mice and treatments.** 8–10-week-old mice were anesthetised with isoflurane and infected intranasally with $3x10^4$ $TCID_{50}$ IAV X-31 in a volume of 30 µl. Weight and health status were monitored daily. Whole lungs or bronchoalveolar lavage (BAL), blood, spleen and lymph nodes were collected at various time points post-infection. For depletion experiments, 100 µg anti-IL-1β antibody or isotype control was administered intraperitoneally in a volume of 200 µl at day 1 and day 2 post-infection. 100 µg anti-IL-17A, 150 µg anti-Ly6G or isotype control antibody (all antibodies BioXcell) were administered intraperitoneally in a volume of 200 µl at -1 day, -1 hour prior- and +1 day post-infection.

**Viral load assessment.** Virus was quantified in infected lungs by qPCR for the Influenza A Matrix gene M1. Whole lungs were collected in RNAlater solution (Invitrogen) and RNA was isolated using TriReagent-Chloroform extraction followed by isopropanol precipitation (all chemicals from Sigma-Aldrich). The TaqMan RNA-to-CT *1-Step* Kit in combination with the PCRmax Human Influenza A virus M1 kit was used quantify M1 copy number.

**Histology.** Whole lungs were harvested at various time points post-infection, fixed overnight in 4% paraformaldehyde (PFA), embedded in paraffin, sectioned and stained with hematoxylin and eosin. Images were acquired using the NanoZoomer slide scanner system (Hamamatsu). Histopathological analysis was performed, blinded to groupings and genotype for various parameters as follows: bronchiolitis classified as damage to airway epithelial cells, necrotic bodies, or denudation of airway epithelial lining; alveolitis classified as damaged alveolar epithelial cells or denuded epithelial lining; bronchiole inflammation classified as bronchioles filled with inflammatory cells; haemorrhage classified as presence of erythrocytes in the alveolar airspace, damaged capillaries, and haemorrhagic effusions in the damaged areas; interstitial inflammation classified as inflammation in the alveoli or thickening of the alveolar interstitium; endothelial damage classified as necrotic endothelium present within small blood vessels or capillaries. Damage severity was scored on a 4-point scale: 0 indicates none or very minor; 1 indicates mild; 2 indicates intermediate; 3 indicates moderately severe; and 4 indicates severe and widespread.

**BAL analysis.** Bronchoalveolar lavage (BAL) of the whole lung was performed with 0.5 mL saline via a tracheal cannula. Samples were centrifuged at 3000 rpm for 10min at 4°C and supernatant was analysed for IFNγ, IL-17A, IL-17F, IL-1β, IL-6, KC, MCP-1, TNFα using Cytometric bead array (CBA) (BD Bioscience) as well as proteome profiler mouse XL cytokine array (R&D) according to the manufacturer's instructions. Sandwich ELISA kits were used for the detection of mouse IL-1β, IL-17A, IFNα4 (all Invitrogen), IFNβ (PBL), IFNλ (Biotechne), PGE2 (LSBio) and TNFα (Stemcell).

**Cell culture.** Mouse neutrophils were isolated from the bone marrow of C57Bl/6J and Nox2-/- mice using the EasySep Mouse Neutrophil Enrichment kit (Stemcell technologies) according to the manufacturer's instructions. Mouse γδ T cells were isolated from lymphoid tissues (spleen and lymph nodes) harvested from C57Bl/6J mice

using the EasySep Release Mouse APC Positive Selection kit (Stemcell technologies) and 2 µg/ml anti-γδTCR antibody (GL3, Biolegend).
Human neutrophils and peripheral blood mononuclear cells (PBMC) layers were freshly isolated over Histopaque-1119 gradient followed by a discontinuous Percoll gradient for neutrophil isolation as described previously by Aga et al. Human γδ T cells were isolated from PBMCs using the EasySep human γδ T cell Isolation kit (Stemcell technologies) according to the manufacturer's instructions.
Mouse and human neutrophils alone or in co-culture with γδ T cells were plated in Hank's balanced salt solution plus $Ca^{2+}$ and $Mg^{2+}$ supplemented with 3% human plasma or 10% fetal calf serum (FCS) for mouse neutrophils, respectively. Cells were pretreated for 1 hour with 10 µM NADPH oxidase inhibitor diphenyleneiodonium (DPI, Cambridge Bioscience) or 300 µM Nox2 inhibitor Apocynin (Merck). Mouse cultures were infected with IAV strain PR8 at MOI 0.2. Human cultures were stimulated with 1 µg/ml poly(I:C) or infected with SARS-CoV-2 England 02 at MOI 0.2.

**Flow cytometry.** Mouse tissues were processed into single-cell suspensions by passing through 70µm cell strainers. Lung tissue was digested in 0.4 mg/ml Liberase TL (Roche) for 45 minutes at 37°C with shaking. After erythrocyte lysis with Ammonium-chloride-potassium (ACK) buffer, cells were washed, filtered and resuspended in Iscove's modified Dulbecco's medium (IMDM) supplemented with 10% FCS.
For cell surface staining, $1x10^6$ cells per sample were incubated with 2.5 - 5 µg/ml antibodies for 15 minutes at room temperature, washed, and fixed in Fixation buffer (Biolegend) for 20 minutes at 4°C.
For intracellular cytokine staining, $2x10^6$ cells were incubated at 37°C for 5 hours with 1x cell activation cocktail (Biolegend). Protein transport blockers 1x Brefeldin and 1x Monensin (both Biolegend) were added for the last 3 hours of incubation. Cells were fixed and permeabilised in BD Cytoperm/Cytofix buffer for 20 minutes at 4°C, blocked for 10 minutes with 10 µg/mL TruStain FcX and stained with 5 - 10 µg/ml antibody for 1 hour at room temperature in the dark.
Dead cells were excluded with the LIVE/DEAD™ Fixable Aqua Dead Cell Stain Kit (Invitrogen). Samples were filtered and resuspended in PBS supplemented 5% FCS (FACS buffer) for acquisition on a Fortessa cell analyser (BD Biosciences) and analysed using FlowJo v10.9 (BD Biosciences).
For flow cytometric imaging, $1x10^6$ cells per sample were incubated with 5 µM CellROX™ Green Reagent, for oxidative stress detection (Invitrogen) for 30 minutes at 37°C, washed with FACS buffer, and stained with 2.5 - 5 µg/mL antibodies for 15 minutes at 4°C protected from light. Samples were washed, resuspended in FACS buffer, and 4',6-diamidino-2-phenylindole (DAPI) was added immediately before acquisition on an ImageStream X analyser (Amnis Corporation). Imagestream data was acquired using INSPIRE and analysed using IDEAS software (both Amnis Corporation).

**Immunofluorescence staining and confocal microscopy of fixed cells and tissue.** Mouse lungs were collected 1, 2, 3 and 5 days post-infection, fixed overnight in 4% paraformaldehyde, and dehydrated in 20% sucrose for 72 hours. After embedding in OCT (Optimum Cutting Temperature compound, VWR) lungs were frozen on dry ice and sectioned at mid-levels on a Leica CM3050 S Cryostat.

$1x10^5$ human or mouse neutrophils were plated on poly-D-lysine-coated glass coverslips 1 hour before stimulation. After various time points cells were fixed in 2% Image-iT™ fixative solution (Invitrogen) for 20 minutes at 37°C and washed 3 times in phosphate-buffered saline (PBS).

Fixed cells and lung sections were permeabilised with 0.5% Triton-X 100 in PBS for 2 minutes, blocked with animal-free blocking diluent (2B Scientific) for 1 hour and incubated with anti-IAV nucleoprotein (Invitrogen), anti-mouse Ly6G-Alexa Fluor 647 (clone 1A8), anti-hamster γδTCR (both Biolegend), or anti-human/mouse p47-phox (Santa Cruz), p47-phox and anti-p67-phox (both Mybiosource) antibodies. Primary antibodies were detected with Alexa Fluor-conjugated secondary antibodies. Cells and tissue sections were mounted in ProLong™ Diamond containing DAPI (Invitrogen) and imaged with a Nikon A1R confocal microscope. Images were analysed with ImageJ software.

Colocalisation of p47-phox/p67-phox and p40-phox/p67-phox was assessed using a Python (v3.9) script in JupyterLab (v3.6.7) based on calculation of Pearson's correlation coefficient available on GitHub (Repository: AnniWarn/Colocalisation).

**DHE staining.** For *in situ* ROS detection mouse lungs were collected 18 hours post-infection in PBS at room temperature. Immediately, lungs were embedded in OCT medium (VWR), frozen on dry ice and sectioned at 30 μm using a Leica cryostat. Sections were mounted on Superfrost Plus glass slides and incubated with 10 μM dihydroethidium (DHE) for 30 minutes at 37°C in the dark, then washed 3 times for 5 minutes each in PBS. Coverslips were added with ProLong™ Diamond containing DAPI (all from Invitrogen) and immediately imaged with identical settings on a Nikon A1R confocal microscope.

For assessment of *in vivo* superoxide production by flow cytometry, BAL was collected in ice-cold PBS supplemented with 10% FCS and centrifuged at 300xg for 6 minutes at 4°C. Cell pellets were resuspended in PBS and $1x10^6$ cells stained with 100 μM DHE for 45 minutes at 37°C in the dark. Cells were washed twice with FACS buffer, fixed in Fixation buffer (Biolegend) for 20 minutes at 4°C, and resuspended in FACS buffer for analysis on a Fortessa cell analyser (BD Biosciences).

**Oxyburst assay.** For assessment of *ex vivo* ROS production, $1x10^6$ isolated human neutrophils were plated in 96-well plates and left to settle for 1 hour. Cells were incubated with 10 μM OxyBURST® Green H2DCFDA (Invitrogen) in Krebs-Ringer PBS (KRP buffer), which contains phosphate-buffered saline pH 7.4 with 1 mM $Ca^{2+}$ , 1.5 mM $Mg^{2+}$

and 5.5 mM glucose. Samples were stimulated and acquired at 0, 10 and 30 minutes on a Fortessa cell analyser (BD Biosciences).

**Luminol assay.** For plate-based *ex vivo* ROS detection, $1x10^6$ isolated human neutrophils were plated in white Nunclon 96-well plates and left to settle for 1 hour at 37°C. Total ROS production was measured by adding 100 µM cell-permeable luminol, extracellular ROS were assessed by adding 100 µM cell-impermeable isoluminol both in the presence of 1.2 U/ml horseradish peroxidase (all chemicals Sigma-Aldrich). Chemiluminescence was measured immediately after stimulation every 30 seconds over 2 hours on a Spectramax plate reader (Molecular devices).

**Quantitative real-time RT-PCR.** Total RNA from mouse lungs stabilised in RNAlater solution (Invitrogen) as well as mouse and human neutrophils was isolated using TriReagent-Chloroform extraction followed by isopropanol precipitation (all chemicals from Sigma-Aldrich). For analysis of interferon mRNA levels samples were additionally purified using the RNeasy kit (Qiagen). Subsequently, 1 µg RNA were reverse transcribed to generate cDNA with the Transcriptor First strand cDNA synthesis kit from Roche using anchored-oligo$(dT)_{18}$ primer. Gene expression was measured in triplicates using TaqMan Universal PCR Master Mix with gene-specific primers and probes on a QuantStudio™ 7 Real-Time PCR machine (all from Applied Biosystems). The cycling-threshold (CT) for each gene was measured and normalised to that of the housekeeping gene (HPRT1 or B2M). The relative gene expression was calculated by the change-in-cycling-threshold (ΔΔCT) method.

**Immunoblot analysis.** $1x10^6$ cells were seeded in 6-well plates and left to settle for 1 hour. Cells were lysed 5 hours post-stimulation in Radio-Immunoprecipitation Assay (RIPA) buffer containing 10 mM Tris-HCl pH 8.0, 1 mM ethylenediaminetetraacetic acid (EDTA), 0.5 mM ethyleneglycoltetraacetic acid (EGTA), 1% Triton X-100, 0.1% Sodium Deoxycholate, 0.1% Sodium dodecyl sulfate (SDS), 140 mM Sodium chloride freshly supplemented with protease inhibitor cocktail 1x complete Mini (Roche), 1 µM Epoxomicin (Generon) and 10 mM N-Ethylmaleimide (Cambridge Bioscience). Lysates were centrifuged at 14000 rpm for 15 minutes to separate cell debris. Protein concentration was determined using the Bicinchoninic acid (BCA) protein assay kit (Pierce). 25 µg of total protein lysate was diluted in SDS loading buffer, boiled at 95°C for 5 minutes, loaded onto NuPAGE 4-12% BisTris gel (Invitrogen) and transferred onto nitrocellulose membranes. Membranes were incubated with 4 µg/ml anti-MPO (R&D) and 1 µg/ml anti-NFkB p105/p50 antibody (Abcam) at 4°C overnight. Secondary anti-IgG antibodies were coupled to HRP (horseradish peroxidase; Thermo Scientific) and visualised by ECL reaction using a ChemiDoc XRS imaging system (Bio-Rad).

**Subcellular fractionation and p50 binding assay.** $5\times10^6$ neutrophils were seeded in 10 cm cell culture dishes. 45 minutes post-stimulation cells were lysed using the Nuclear

Extraction Kit (Cambridge Bioscience) according to the manufacturer's instructions. Nuclear lysates containing 10 µg protein were used in a TransAM® NFκB p50 transcription factor assay (Active Motif).

**Enzyme linked Immunosorbent assay.** $4x10^5$ neutrophils human or mouse neutrophils were seeded in a 96-well U-bottom plates. Cell culture supernatant was collected 5 hours post-stimulation. In coculture experiments, murine γδ T cells and neutrophils were combined in a 4:1 ratio. For cell interaction studies, γδ T cells were seeded in the bottom compartment of a Transwell plate. To block IL-1β signalling, 5 µg/ml anti-IL-1β antibody (BioXcell) was added. Supernatants were analysed using human IL-1β, mouse IL-1β and mouse IL-17A ELISA kits (all from Invitrogen) according to the manufacturer's instructions.

**Quantification and statistical analysis.** GraphPad Prism 10 was used for data presentation and statistical analysis. Statistical parameters, used tests, and exact number of replicates (n) is given in the figure legends. Data are presented as mean and SEM unless indicated otherwise. *P* values of less than 0.05 were considered significant.
