## Supplemental figures for "Neutrophil NADPH oxidase breaks the inflammatory IL-1β/IL-17A circuit to enhance pathogen clearance during respiratory virus infections"

### Supplemental information

**Supplemental figure S1.** Verification of neutrophil-specific Nox2 deletion.

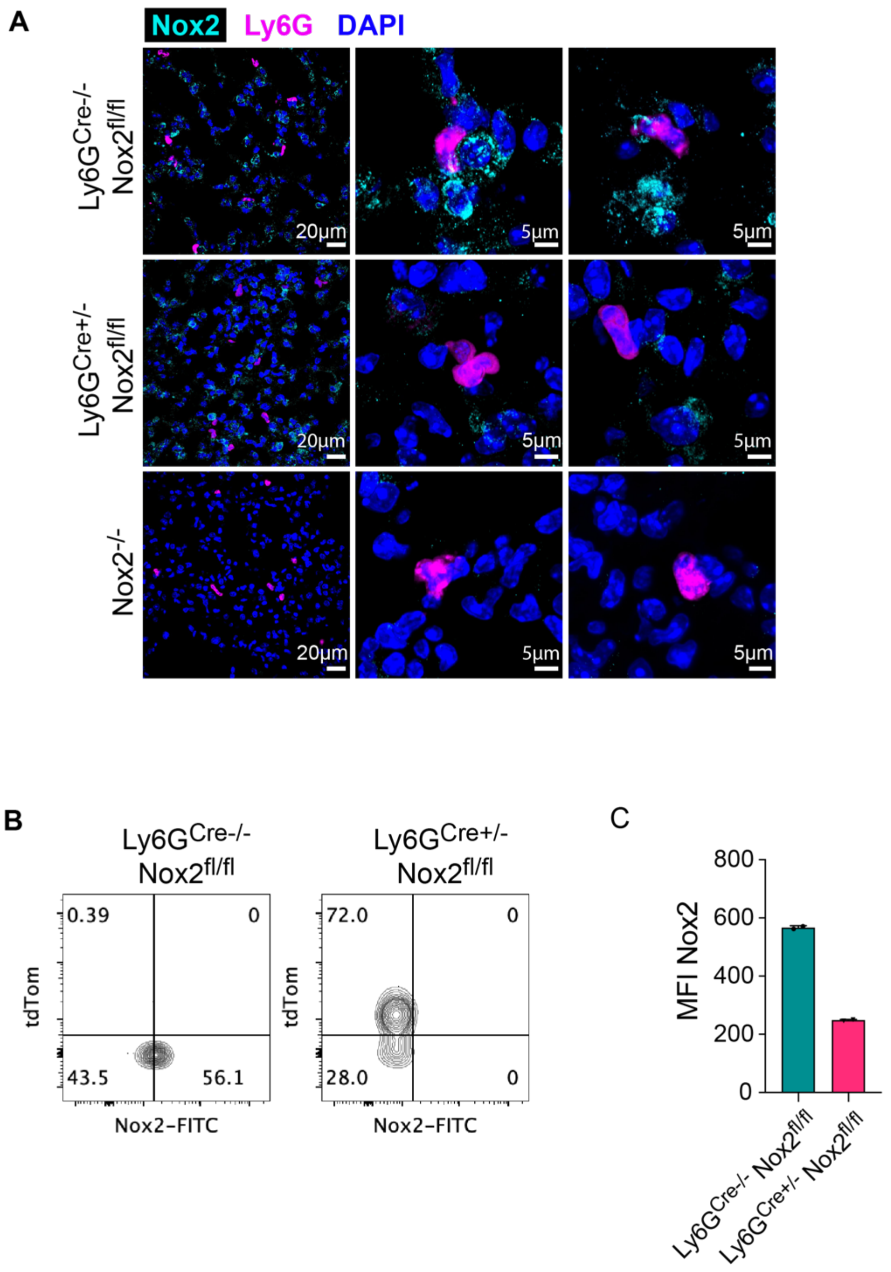

**Supplemental figure S1.** Verification of neutrophil-specific Nox2 deletion.

A) Representative micrographs of lung sections of naïve *Ly6G-Cre*$^{+/-}$, *Ly6G-Cre*$^{+/-}$ *Nox2*$^{fl/fl}$ and *Nox2*$^{-/-}$ mice stained for neutrophils (Ly6G$^{tdTom}$, magenta), Nox2 (gp19phox, cyan) and nuclei (DAPI, blue). Overview at 20x magnification (left panels). Scale bar 20 μm. Magnification of single cells at 60x (right panels). Scale bar 5 μm.

B) Representative FACS plots of tdTom$^{+}$ versus Nox2$^{+}$ blood-isolated neutrophils of naïve *Ly6G-Cre*$^{-/-}$ *Nox2*$^{fl/fl}$ and *Ly6G-Cre*$^{+/-}$ *Nox2*$^{fl/fl}$ gated as Live CD45$^{+}$, CD11b$^{+}$ and Ly6G$^{+}$.

C) Nox2 mean fluorescence intensity (MFI) of neutrophils.

**Supplemental figure S2.** Virus burden, weight loss and histopathology.

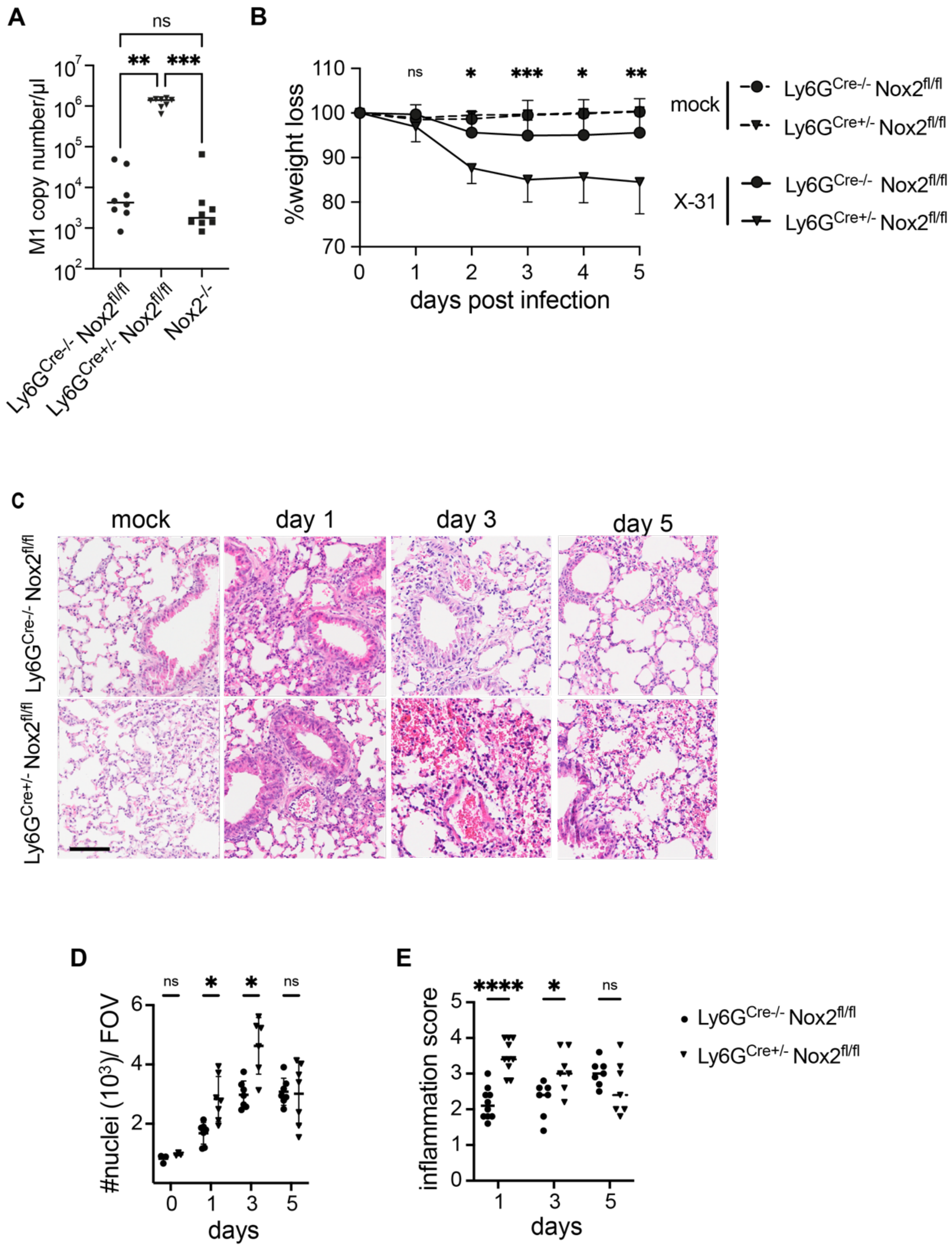

**Supplemental figure S2.** Virus burden, weight loss and histopathology.

A) RT-qPCR of influenza matrix protein M1 in whole lungs 3 days post-infection with IAV strain X-31 at $3x10^4$ $TCID_{50}$ in *Ly6G-Cre$^{-/-}$ Nox2$^{fl/fl}$*, *Ly6G-Cre$^{+/-}$ Nox2$^{fl/fl}$* and *Nox2$^{-/-}$* mice (n = 8).

B) Weight loss monitored over 5 days post-infection with IAV strain X-31 at $3x10^4$ $TCID_{50}$ in *Ly6G-Cre$^{-/-}$ Nox2$^{fl/fl}$* and *Ly6G-Cre$^{+/-}$ Nox2$^{fl/fl}$* mice (n = 8).

C) Representative micrographs of H&E-stained lung sections at indicated time points after infection with $3x10^4$ $TCID_{50}$ X-31 in *Ly6G-Cre$^{+/-}$* and *Ly6G-Cre$^{+/-}$ Nox2$^{fl/fl}$* mice. Scale bar 100 µm.

D) Number of nuclei per field of view at indicated time points after infection with $3x10^4$ $TCID_{50}$ X-31.

E - F) Scoring of inflammation for bronchiolitis (E) and alveolitis (F) at indicated time points after infection with $3x10^4$ $TCID_{50}$ X-31. The degree of inflammation was graded as follows: 0, normal; 1, mild; 2, moderate; 3, severe; and 4, very severe inflammation.

Bars represent mean ± SEM of 2 – 3 independent experiments (A - F). p values are indicated, *<0.05, **<0.01, ***<0.001, ****<0.0001, ns not significant. p values were determined using Kruskal-Wallis test and Dunn's multiple comparisons post-test (A) two-way ANOVA and Sidak's post-test (B, D, E, F).

**Supplemental figure S3.** Immune cell infiltration and cytokine array.

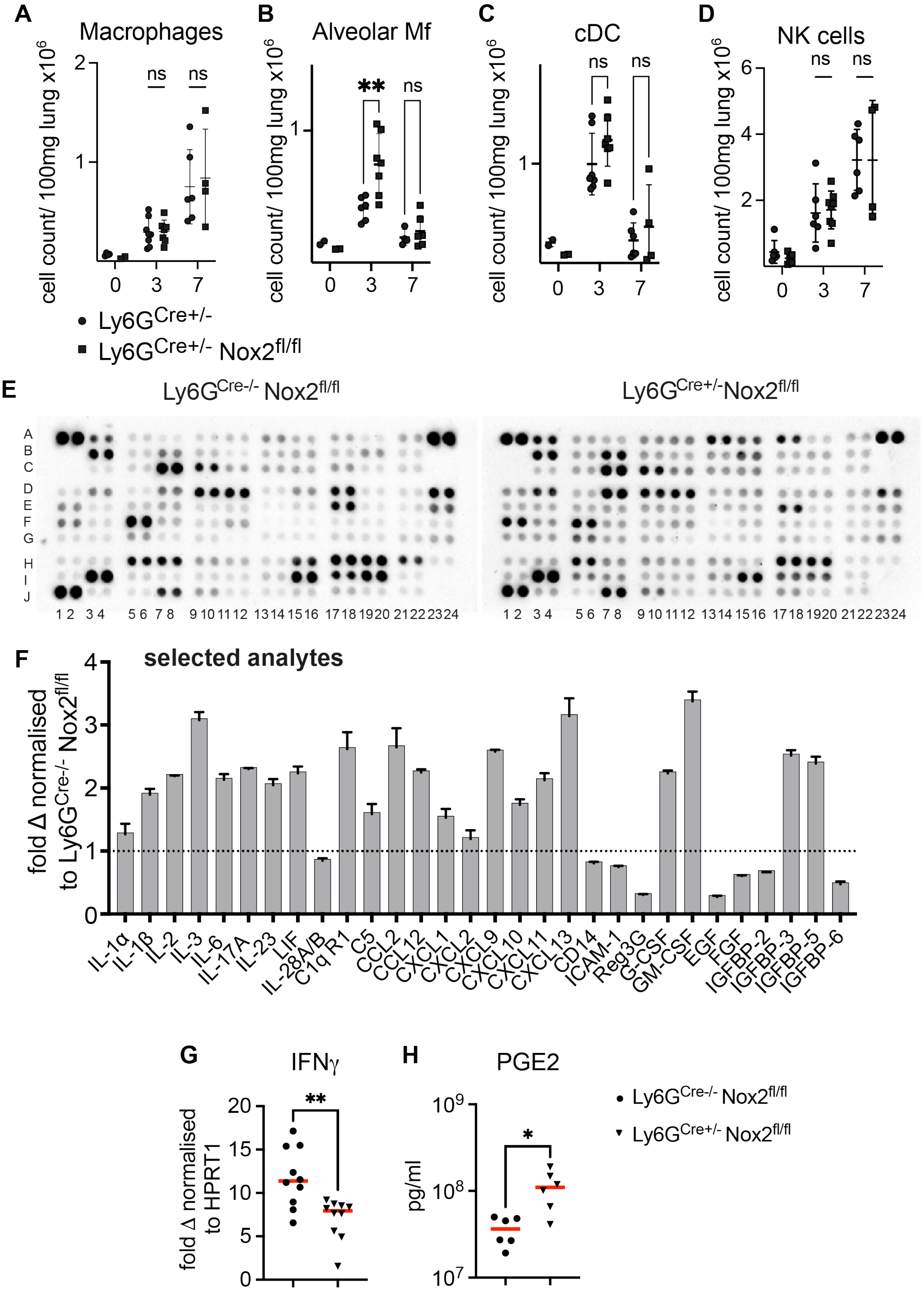

**Supplemental figure S3.** Immune cell infiltration and cytokine array.

*Ly6G-Cre*$^{+/-}$, *Ly6G-Cre*$^{-/-}$ *Nox2*$^{fl/fl}$ and *Ly6G-Cre*$^{+/-}$ *Nox2*$^{fl/fl}$ mice were infected with $3x10^4$ $TCID_{50}$ IAV strain X-31 or PBS (mock) in 30 µl intranasally.

A – D) Number of mature macrophages gated as Live $CD45^+$, $CD11b^+$, $F4/80^+$ and $MHCII^+$ (A), Alveolar macrophages gated as $CD11c^+$ and Siglec-$F^+$ (B), conventional dendritic cells gated as $CD11c^+$, $CD103^+$ and $MHCII^+$ (C) and NK cells gated as $CD3e^-$ and $NK1.1^+$ (D) per 100 mg lung tissue 3 and 7 days post-infection (n = 2 - 4 mock, n = 4 - 8 X-31).

E – F) Membrane-based cytokine and chemokine array (proteome profiler by R&D) on bronchoalveolar lavage of *Ly6G-Cre*$^{-/-}$ *Nox2*$^{fl/fl}$ and *Ly6G-Cre*$^{+/-}$ *Nox2*$^{fl/fl}$ mice infected with IAV strain X-31 at $3x10^4$ $TCID_{50}$ for 3 days. Representative nitrocellulose membranes for one animal of each group. Shown are the coordinates of 111 different analytes. Each cytokine/ chemokine is spotted in duplicate (E). Quantitated in (F) as mean pixel density of selected interleukines and chemokines from the array calculated as fold change from infected *Ly6G-Cre*$^{-/-}$ *Nox2*$^{fl/fl}$ animals (n = 5).

G) IFNγ mRNA expression in fold change from naïve normalised to HPRT1 in lung tissue of *Ly6G-Cre*$^{-/-}$ *Nox2*$^{fl/fl}$ and *Ly6G-Cre*$^{+/-}$ *Nox2*$^{fl/fl}$ animals 3 days post-infection was quantified by RT-qPCR (n = 2 mock, n = 10 X-31).

H) Prostaglandin E2 (PGE2) protein in bronchoalveolar lavage of *Ly6G-Cre*$^{-/-}$ *Nox2*$^{fl/fl}$ and *Ly6G-Cre*$^{+/-}$ *Nox2*$^{fl/fl}$ animals 3 days post-infection was quantified by ELISA (n = 6 X-31).

Bars represent mean ± SEM of 2 – 3 independent experiments (A – D, F). p values are indicated, *<0.05, **<0.01, ***<0.001, ****<0.0001, ns not significant. p values were determined using two-way ANOVA and Sidak's post-test (A - D) and (F).

**Supplemental figure S4.** Cellular cytokine production.

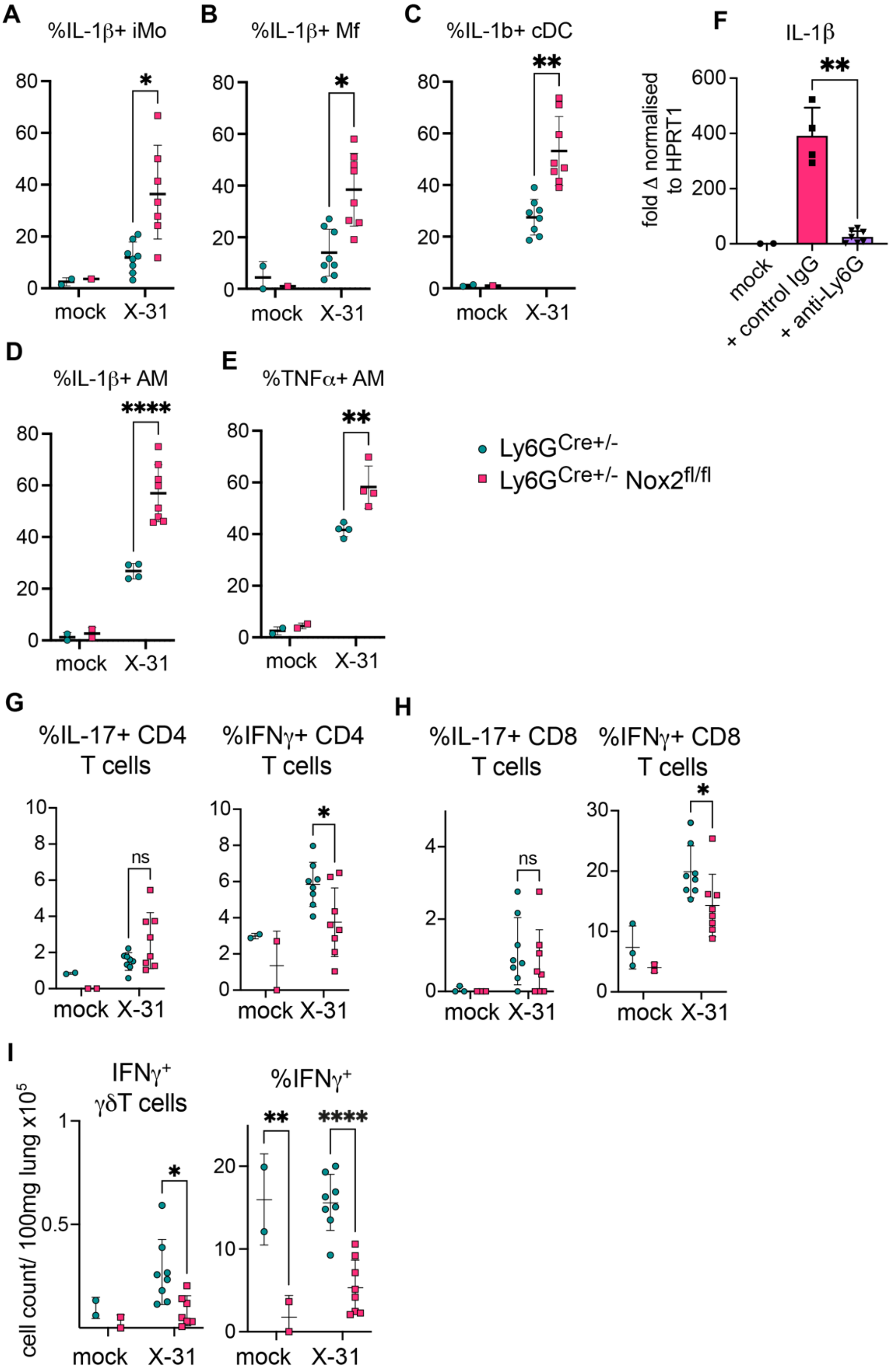

**Supplemental figure S4.** Cellular cytokine production.

*Ly6G-Cre*$^{+/-}$ and *Ly6G-Cre*$^{+/-}$ *Nox2*$^{fl/fl}$ mice were infected with $3x10^4$ $TCID_{50}$ IAV strain X-31 or PBS (mock) in 30 µl intranasally.

A - E) Frequency of IL-1β$^+$ inflammatory monocytes gated as Live CD45$^+$, CD11b$^{hi}$, Ly6G$^-$, Ly6C$^{hi}$ and MHCII$^+$ (A), macrophages gated as CD11b$^+$, F4/80$^+$ and MHCII$^+$ (B), conventional dendritic cells gated as CD11c$^+$, CD103$^+$ and MHCII$^+$ (C), Alveolar macrophages (AM) gated as CD11c$^+$ and Siglec-F$^+$ (D) and TNFα$^+$ AM (E) at 3 days post-infection (n = 2 mock, n = 4 – 8 X-31).

F) Nox2-/- nice were treated with 100 µg anti-Ly6G or control IgG antibody in 200 µl intraperitoneally at -1 day, -1 hour prior- and +1 day post-infection with IAV strain X-31 at $3x10^4$ $TCID_{50}$ or PBS (mock) in 30 µl intranasally. mRNA expression of IL-1β as fold change from naïve normalised to HPRT1 assessed by RTqPCR in lung tissue 3 days post-infection (n = 2 mock, n = 4 – 7 X-31).

G - H) Frequency of IL-17$^+$ and IFNγ$^+$ CD4$^+$ (G) and CD8$^+$ (H) T cells gated as Live CD45$^+$, CD3e$^+$ and γδTCR$^-$ at 3 days post-infection (n = 2 - 3 mock, n = 8 X-31).

I) Count per 100 mg lung tissue (left panel) and frequency (right panel) of IFNγ$^+$ γδ T cells (n = 2 mock, n = 8 X-31).

**Supplemental figure S5.** Proliferation, apoptosis and oxidative stress.

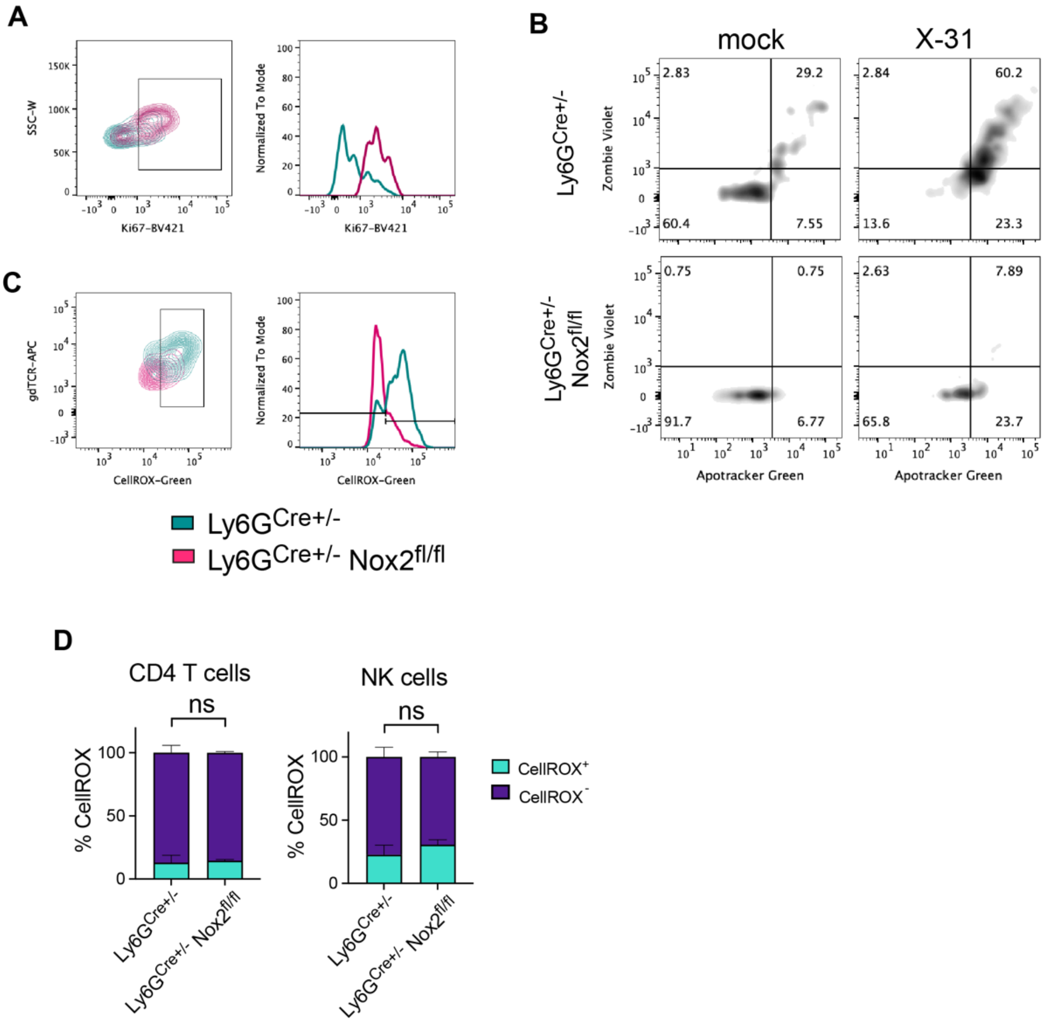

**Supplemental figure S5.** Proliferation, apoptosis and oxidative stress.

*Ly6G-Cre*$^{+/-}$ and *Ly6G-Cre*$^{+/-}$ *Nox2*$^{fl/fl}$ mice were infected with $3x10^4$ $TCID_{50}$ IAV strain X-31 or PBS (mock) in 30 µl intranasally.

A) Representative FACS (left panel) and histogram (right panel) plot of Ki67$^+$ γδ T cells in lungs at 3 days post-infection.

B) Representative FACS plots of ZombieViolet$^+$ and Apotracker$^+$ γδ T cells in lungs at 3 days post-infection.

C) Representative FACS (left panel) and histogram (right panel) plot of CellROX$^+$ γδ T cells in lungs at 3 days post-infection.

D) Frequency of CellROX$^+$ CD4 T cells (left panel, n = 3) and NK cells (right panel, n = 3) in lungs at 3 days post-infection.

**Supplemental figure S6.** Isolation of mouse neutrophils and γδ T cells.

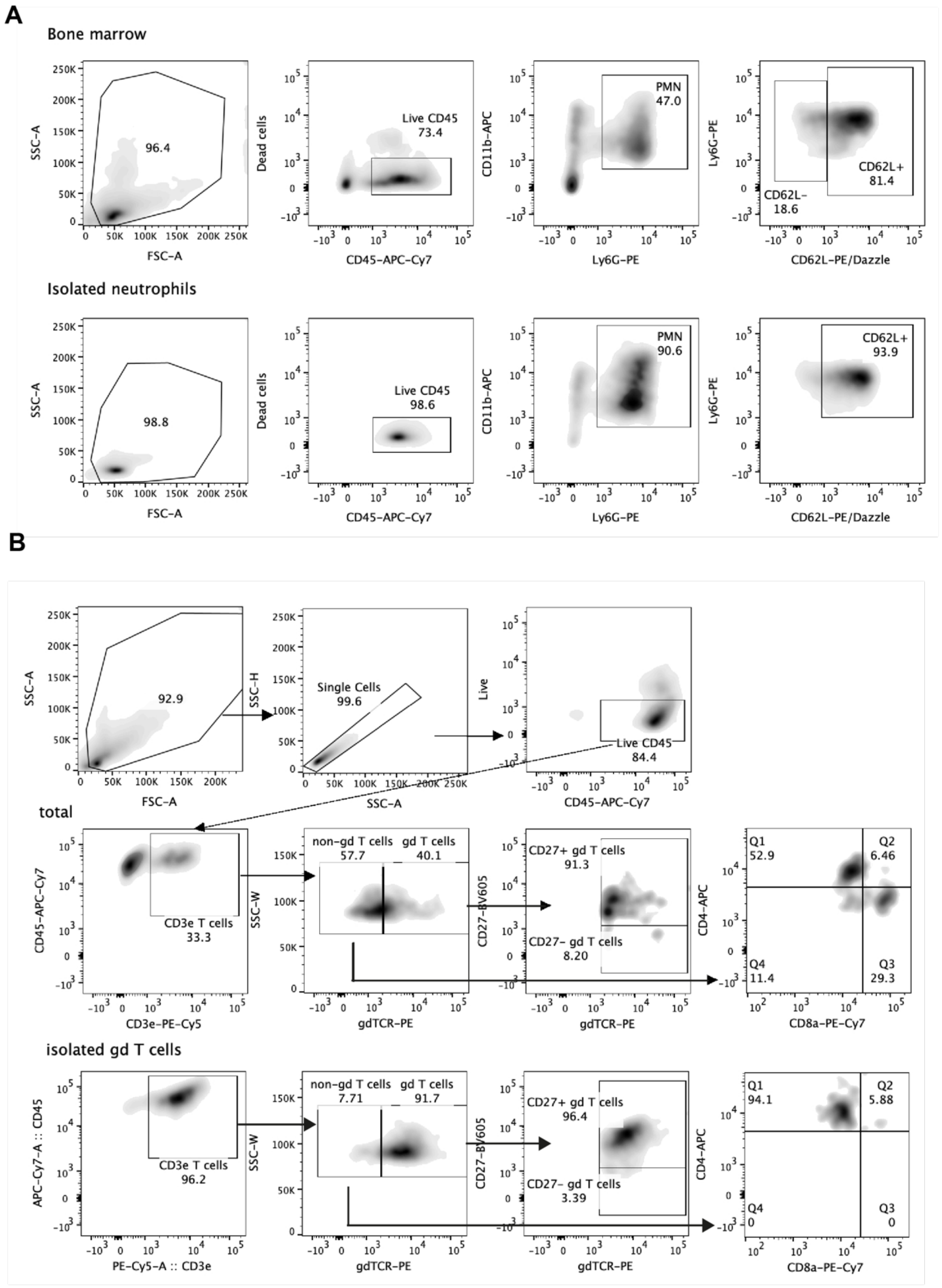

**Supplemental figure S6.** Isolation of mouse neutrophils and γδ T cells.

Neutrophils were isolated from bone marrow and γδ T cells from spleen and lymph nodes of naïve C57Bl/6 and *Nox2*$^{-/-}$ mice.

(A) Gating strategy and representative FACS plots of neutrophils before (upper panels) and after isolation (lower panels) gated as Live CD45$^{+}$, CD11b$^{+}$, Ly6G$^{+}$ and CD62L$^{+}$.

(B) Gating strategy and representative FACS plots of γδ T cells before (upper panels) and after isolation (lower panels) gated as Live CD45$^{+}$, CD3e$^{+}$, γδTCR$^{+}$, CD4$^{-}$ and CD8$^{-}$.

**Supplemental figure S7.** Mouse neutrophil and γδ T cell co-cultures.

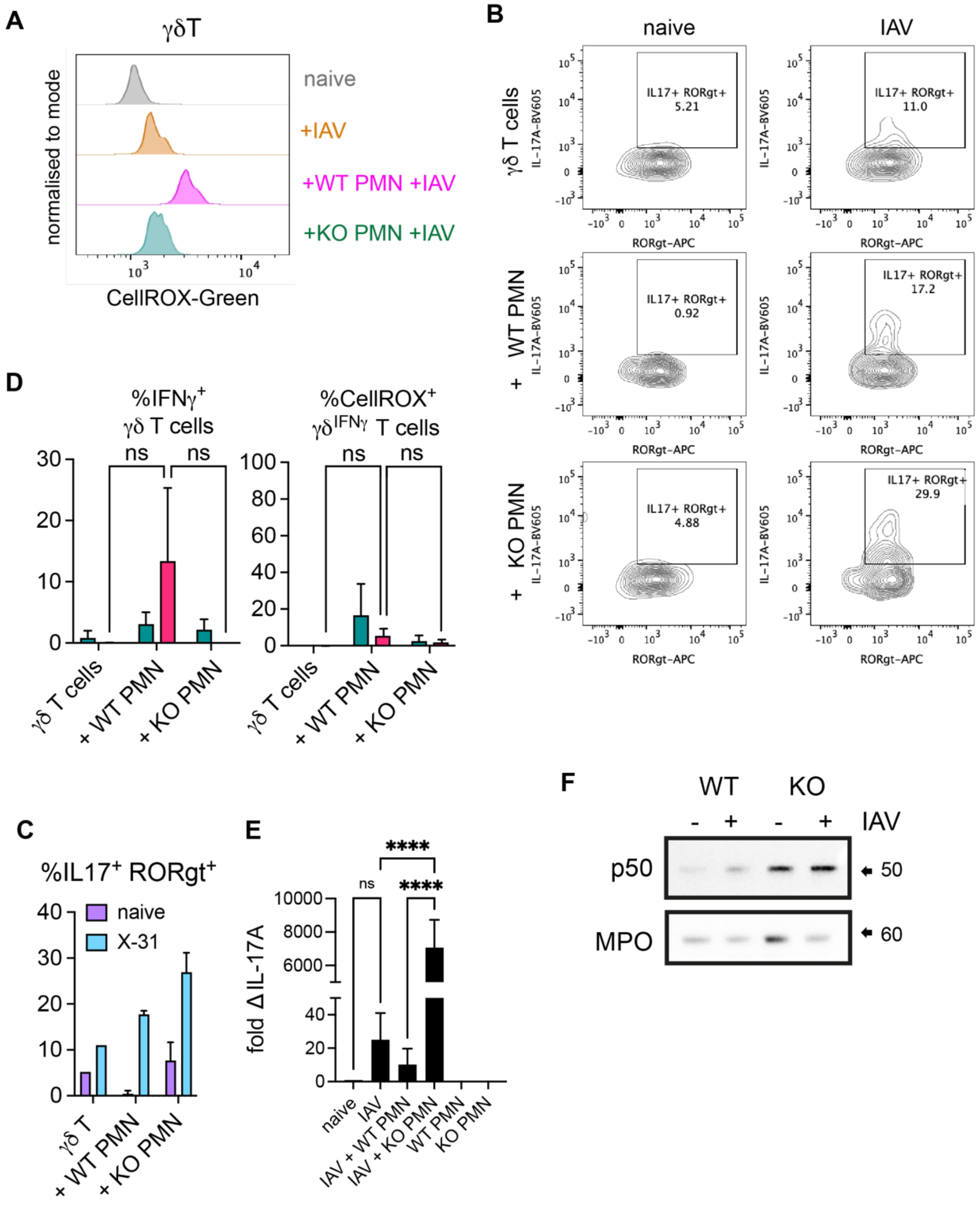

**Supplemental figure S7.** Mouse neutrophil and γδ T cell co-cultures.

Isolated γδ T cells gated as Live $CD45^+$, $CD3e^+$, $CD4^-$ and $γδTCR^+$ were co-cultured *ex vivo* with WT (C57Bl/6) or KO (*Nox2*$^{-/-}$) neutrophils gated as Live $CD45^+$, $CD11b^+$ and $Ly6G^+$ were treated with PBS or infected with IAV strain PR8 at 0.2 MOI (multiplicity of infection).

A) Representative histogram plot of $CellROX^+$ γδ T cells 2 hours post-infection cultured alone, in contact with WT or KO neutrophils.

B - C) Representative FACS plots (B) and Frequency (C) of IL-$17A^+$/$RORγt^+$ γδ T cells 5 hours post-infection cultured alone, in contact with WT or KO neutrophils (n **=** 3).

D) Frequency of $IFNγ^+$ γδ T cells (left panel) and $CellROX^+$ $γδ^{IFNγ}$ T cells (right panel) 5 hours post-infection cultured alone, in contact with WT or KO neutrophils (n = 4).

E) IL-17A mRNA expression of γδ T cells, WT and KO neutrophils alone, γδ T cells co-cultured with WT or KO neutrophils assessed by RT-qPCR 4 hours post-infection (n = 3).

F) Immunoblot analysis of NFκB subunit p50 and myeloperoxidase (MPO) in lysates of mouse WT and KO neutrophils 5 hours post-infection. Representative of 2 independent experiments.

Bars represent mean ± SEM of 3 - 4 independent experiments (C, D). Bars represent mean ± SEM of technical replicates representative of 2 independent experiments (E). p values are indicated, ****<0.0001, ns not significant. p values were determined using two-way ANOVA with Tukey's post-test (D) and one-way ANOVA with Tukey's post-test (E).

**Supplemental figure S8.** Human neutrophil and γδ T cell co-cultures.

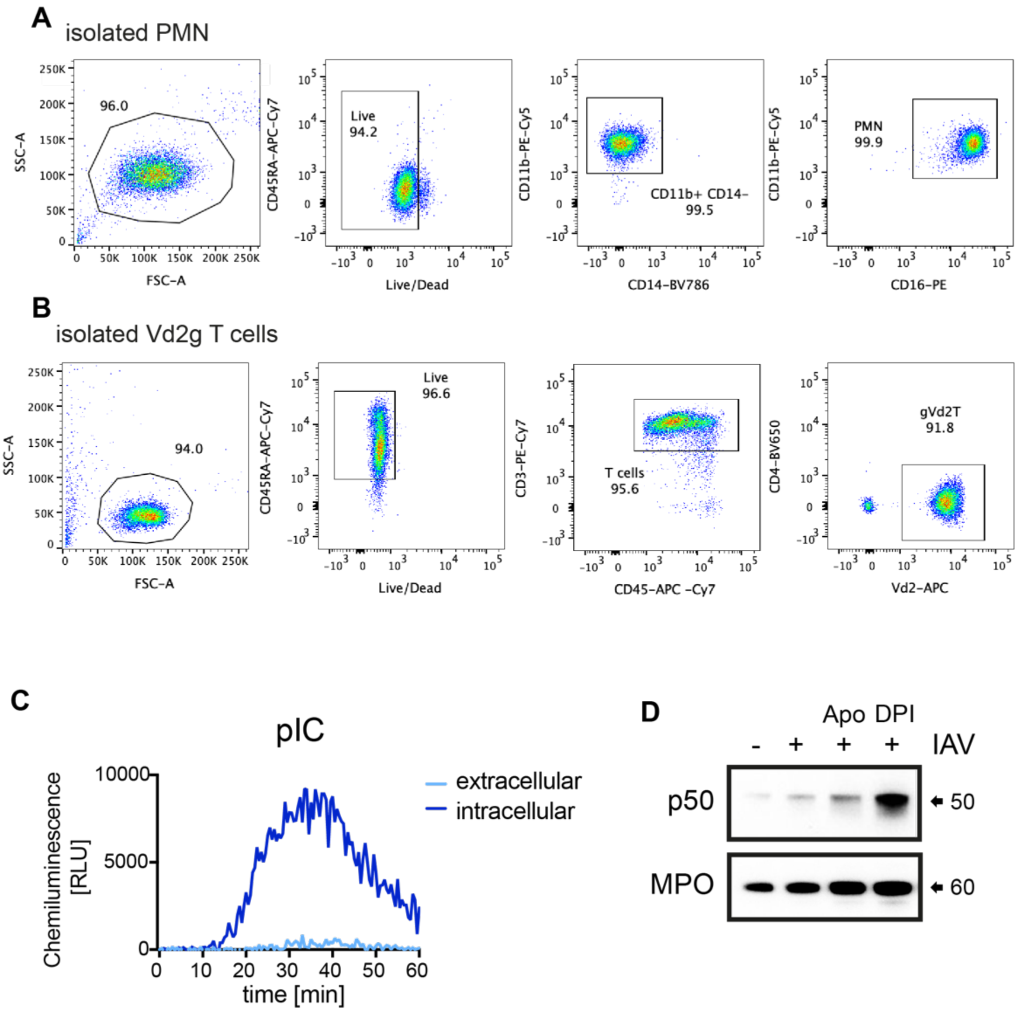

**Supplemental figure S8.** Human neutrophil and γδ T cell co-cultures.

γδ T cells and neutrophils isolated from peripheral blood of healthy volunteers were co-cultured *ex vivo*.

(A) Gating strategy and representative FACS plots of isolated neutrophils gated as Live $CD45RA^-$, $CD11b^+$, $CD14^-$ and $CD16^+$ cells.

(B) Gating strategy and representative FACS plots of isolated γδ T cells gated as Live $CD45^+$, $CD3^+$, $CD4^-$ and $\gamma V\delta 2TCR^+$ cells.

(C) ROS production by human blood-derived neutrophils stimulated with 1 μg/ml poly(I:C) (pIC) detected by chemiluminescence of cell-impermeable isoluminol to detect extracellular ROS or by the cell-permeable luminol to detect intra- and extracellular ROS, in the presence of exogenous horseradish peroxidase to detect superoxide production. Representative of 2 independent experiments.

(D) Immunoblot analysis of NFκB subunit p50 and myeloperoxidase (MPO) in lysates of human neutrophils pre-treated 1 hour with PBS, 300 μM Apocynin (Apo) or 10 μM Diphenyleneiodonium (DPI), stimulated 5 hours with 1 μg/ml pIC. Representative of 2 independent experiments.

Bars represent mean ± SEM of 2 independent experiments (F). p values are indicated, $*<0.05$, ns not significant. p values were determined using two-way ANOVA and Sidak's post-test (F).
